# Machine Learning Approaches in Label-Free Small Extracellular Vesicles Analysis with Surface-Enhanced Raman Scattering (SERS) for Cancer Diagnostics

**DOI:** 10.1101/2024.02.19.581099

**Authors:** Der Vang, Maria S. Kelly, Manisha Sheokand, Manju Sharma, Leyla Esfandiari, Ruxandra I. Dima, Pietro Strobbia

## Abstract

Early diagnosis remains of pivotal importance in reducing patient morbidity and mortality in cancer. To this end, liquid biopsy is emerging as a tool to perform broad cancer screenings. Small extracellular vesicles (sEVs), also called exosomes, found in bodily fluids can serve as important cancer biomarkers in these screenings. Our group has recently developed a label-free electrokinetic microchip to purify sEVs from blood. Herein, we demonstrate the feasibility to integrate this approach with surface-enhanced Raman scattering (SERS) analysis. SERS can be used to characterized extracted sEVs through their vibrational fingerprint that changes depending on the origin of sEVs. While these changes are not easily identified in spectra, they can be modeled with machine learning (ML) approaches. Common ML approaches in the field of spectral analysis use dimensionality reduction method that often function as a black box. To avoid this pitfall, we used Shapley additive explanations (SHAP) is a type of explainable AI (XAI) that bridges ML models and human comprehension by calculating the specific contribution of individual features to a model’s predictions, directly correlating model/decisions with the original data. Using these approaches we demonstrated a proof-of-concept model predictive of cancer from isolated sEVs, integrating the electrokinetic device and SERS. This work explores the use of explainable AI to perform diagnostic analysis on complex SERS data of clinical samples, while reporting interpretable biochemical information.

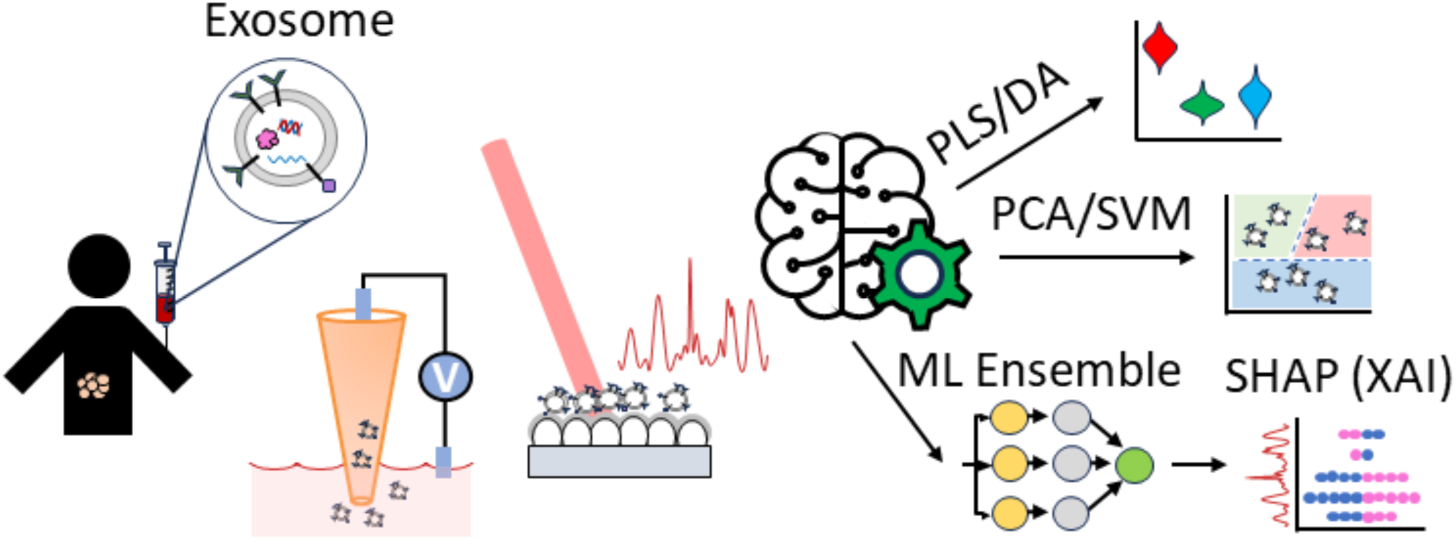

## Introduction

Early diagnosis remains of pivotal importance in reducing patient morbidity and mortality in cancer.^1, 2^ Current practice involves testing with relatively invasive and costly methods only after the onset of symptoms and is further confirmed with pathology analysis of tissue biopsy, the current gold standard for cancer diagnosis.^3–5^ A big issue with this diagnostic paradigm is that it is reactive rather than proactive, which causes diagnosis to arrive after the disease has progressed reducing treatment-positive outcomes. Conversely, non-invasive cancer screening is emerging as a powerful option to provide beneficial early diagnosis, as demonstrated by the popularity of protocols such as Cologuard (limited to colon cancer).^6^ Thereby, there is a strong need for non-invasive methods to detect and monitor cancer.

Liquid biopsy is a non-invasive alternative to tissue biopsy to perform cancer screening.^7–10^ Liquid biopsies consist of the analysis of various bodily fluids (e.g., blood, plasma, saliva, or urine samples). Bodily fluids can be collected non-invasively and contain tumor-related biomarkers. Among those biomarkers, small extracellular vesicles (sEVs), known as exosomes (i.e., nanovesicles of 30-150 nm secreted by various cell types), can be found in bodily liquids. Recently research has shown tumor-derived SEVs carry tumor-specific biomolecules, including RNA, DNA, lipids, and protein from their parent cells.^11^ Because of their biochemical relation with the parent cell, sEVs have been proposed as a promising biomarker for early diagnosis and prognosis prediction for patients, thus, promoting the chances of successful and curated treatments.^12–15^ These sEVs, can be detected and quantified via Western blots, polymerase chain reaction (PCR), enzyme-linked immunosorbent assay (ELISA), and sequencing techniques.^15^ However, these techniques require prior knowledge of specific markers to target and can be time-consuming.

Microfluidic devices based on electrokinetics represent a category of tools designed to isolate sEVs using external electrical forces, eliminating the need for tagging or labeling specific molecules of interest.^17, 18^ The dielectrophoresis (DEP) phenomenon, harnessed in these devices, enables precise isolation based on the dielectric properties and size distribution of sEVs under a non-uniform electric field, offering a rapid and label-free approach. This non-uniform electric field can be generated through the application of an alternating current across an array of electrodes^19, 20^ or by employing an insulator-based dielectrophoretic (iDEP) strategy using obstacles such as micro-pillars in microfluidic channels.^21^ The iDEP approach effectively manipulates biomolecules while maintaining device functionality, despite potential fouling effects on electrode surfaces.^22^ However, the majority of existing devices in this category necessitate high operational voltage (around ∼100 V/cm), posing a risk of denaturing the molecules of interest.^23^ Additionally, despite their promising attributes, these devices come with inherent challenges, including high fabrication costs, the requirement for sample dilution, and the susceptibility of micro- and nano-scale channels to clogging.^24^

Our group has introduced a novel class of iDEP devices designed for the rapid and selective entrapment of nanoparticles based on their size and distinctive dielectric properties. This device, featuring an array of micropipettes, can isolate nanoparticles from small sample volumes by applying a significantly low electric field (∼10 V/cm).^25^ The minute conical pore geometry of the micropipettes enables the isolation of sEVs without the need for sample dilution, allowing downstream analyses with uncompromised yield. Previous demonstrations have showcased the capability of our iDEP device in isolating sEVs from small sample volumes (∼200 μL) of conditioned cell culture media and biofluids from healthy donors within a mere 20 minutes.^26.27^ The device also has the capability to characterize the isolated sEVs based on their unique dielectric properties.^28–30^

The next step is the characterization of sEVs. sEVs from different cells can have very similar biomolecular information (e.g., protein receptors), making targeting and distinguishing cancer-related sEVs markers a challenge.^31^ Using sEVs for clinical diagnostics remains a challenge, and there is a need for a more robust classification of sEVs subpopulations, and sEVs isolation methods.^14^ Raman spectroscopy is a powerful spectroscopic technique that permits the identification of molecules through their vibrational fingerprint. These features can be used for diagnostic purposes by collecting biochemical information from different sEVs. Raman spectra are characterized by low signals making it challenging to detect significant signals from analytes at ultralow concentrations, such as sEVs. To overcome this limitation, the use of plasmonic metal nanomaterial or nanostructures can amplify weak Raman signals via surface-enhanced Raman scattering (SERS). In recent years, SERS has been shown to be able to identify sEV origins and other EVs, by incorporating multivariate analysis and machine learning algorithms as classification tools.^32–41^ These tools can extract small variations in the SERS spectral profile and differentiate the sEVs.

Machine learning (ML) techniques have emerged -as a transformative tool in the field of spectral analysis, where high dimensional datasets composed of hundreds of peak intensities are simplified in a meaningful way to extract patterns that enable rapid and accurate classification.^42–44^ For cancer detection, a model can be built from numerous samples of both healthy and diseased patients to learn the essential relationships between a collection of peaks that can then be applied to samples of new patients for diagnostics.^45, 46^ Therefore, the ability of SERS to examine sEVs coupled with ML can reduce the complexity found within many spectra and can be used for the effective classification of healthy and cancerous sEVs.^46–48^

Building an ML classification model, however, can be a challenging endeavor, and it often yields misleading or inaccurate results when not meticulously tuned. One of the challenges lies in selecting the appropriate algorithm for the specific dataset, as well as balancing the model’s complexity to prevent overfitting or underfitting.^49^ Preprocessing steps with dimensionality reduction techniques, such as partial least squares (PLS) and principal component analysis (PCA), must be carefully executed to avoid removing critical information needed for proper generalization of the model. Furthermore, failure to optimize hyperparameters, such as learning rate or tree depth, is essential for reducing the bias in a model’s predictive ability. Ignoring these aspects can result in a model that appears to perform well during training but fails at generalizing the differences between samples, which can lead to misguided diagnoses for patients.

While well-tuned ML models have shown remarkable capabilities in making accurate predictions, being able to interpret the predictive decisions made by the model can bolster confidence in its generalization and identify features that highly influenced the predictions made. To bridge ML models and human comprehension, explainable AI, specifically Shapley Additive exPlanations (SHAP), can be used to report the contribution of individual features to a model’s predictions.^50^ SHAP is based on cooperative game theory which allows for more consistent weight assignments while also providing both local and global interpretability. Permutation feature importance (PFI) is another common technique that calculates feature’s influence on prediction accuracy but has been shown to be misleading when analyzing complex datasets with highly non-linear relationships.^51^ Local Interpretable Model-Agnostic explanation (LIME) is a popular explainable approach which explains model decisions through creating a separate model of feature perturbations but is limited to only interpreting a single instance.^52^ Therefore, due to the information-rich data produced from SERS analysis, utilizing SHAP can offer a way to gain chemical insight into the classification process and link the classification to the biochemical hypothesis.

Herein, we surveyed various SERS-substrate designs to select the substrate with the highest plasmonic properties for sEV analysis. We performed extensive tuning of multiple models under two main approaches: initial transformation of the data set using dimensionality reduction tools and feature removal after model tuning to reduce the number of dimensions included in the final classification analysis (Figure 1). This present work provides useful information on how to apply and access learning models with SHAP. With SHAP, we extrapolated and identified what Raman bands significantly influenced the classification model. We then applied the optimized classification algorithm approach on SERS spectra collected from sEVs isolated from a kidney cancer patient’s blood via the electrokinetic microchip. These results demonstrate a proof-of-concept model predictive of cancer from isolated sEVs. While these samples do not represent the variability of a pre-clinical study (no patient cohort), each spectrum is a complex mix of sEVs-substrate interactions and heterogenous sEVs composition. We also first demonstrate the integration of two emerging techniques for non-invasive diagnostics (i.e., iDEP and SERS) that can potentially be used in stand-alone devices. As a further demonstration of the feasibility of this integration, we also demonstrated our predictive model using substrates fabricated with colloidal nanoparticles, easily integrated in pre-existing devices. Importantly, our work demonstrates the use of explainable AI to perform diagnostic analysis on complex SERS data of clinical samples. This approach enabled two key advances: -1- optimization of the classification model based on its correlation with specific Raman bands and not just on diagnostic accuracy and -2- classification of patients’ sEVs based on diagnosis while reporting interpretable biochemical information directly correlated to this model.

**Figure 1.**
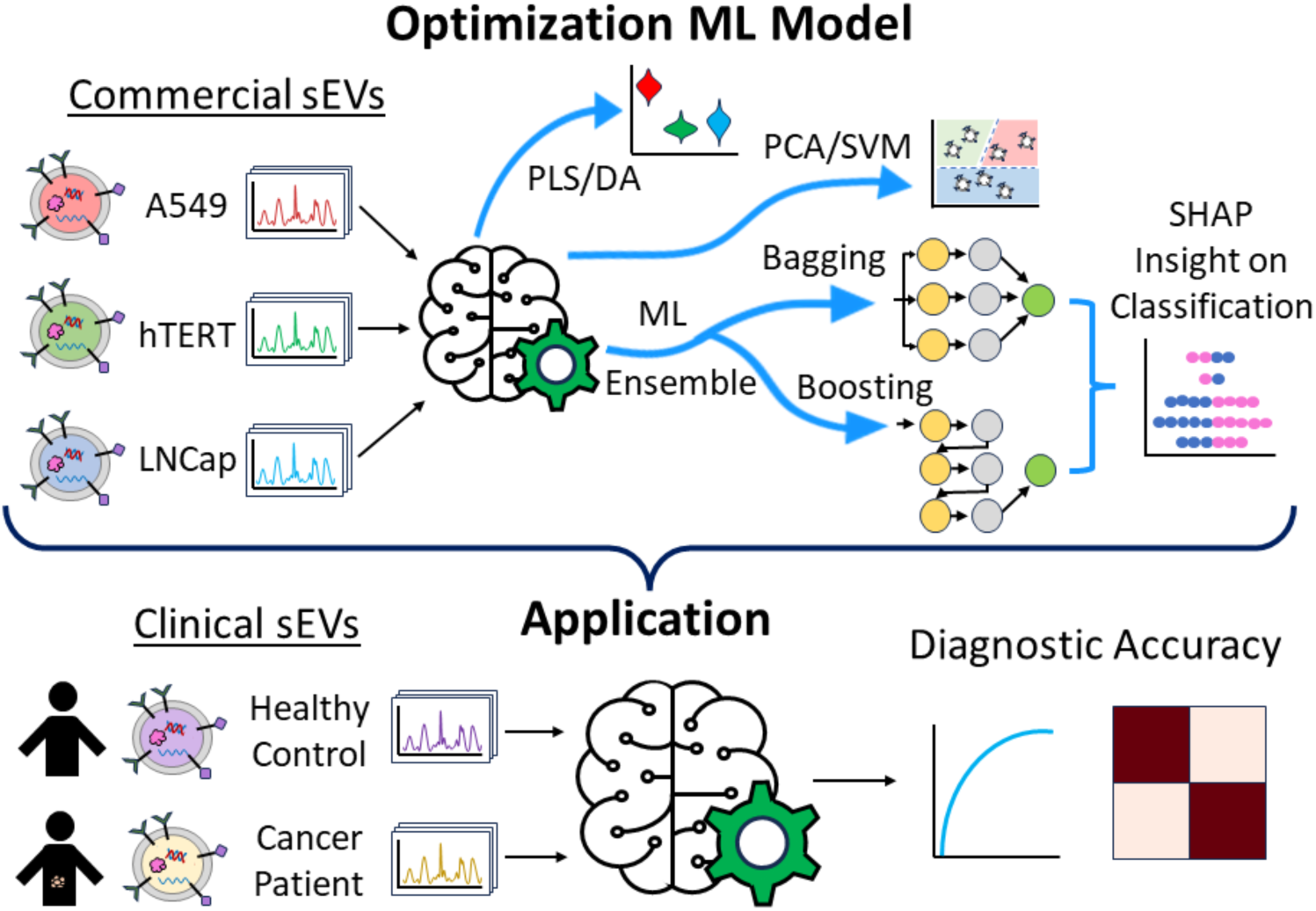
Schematic workflow of machine learning approach of sEVs analysis for cancer diagnosis.

## Experimental

### Materials and chemicals

Hydrogen peroxide (H_2_O_2_), gold(III) chloride trihydrate (HAuCl_4_), silver nitrate (AgNO_3_), sodium citrate tribasic dihydrate (Na_3_Cit), L(+)-ascorbic acid (AA), benzoic acid (4-MBA), and (3-aminopropyl)triethoxysilane 99% (APTES), Liposome Kit: Lipid mixture for the preparation of liposomes were purchased from Sigma-Aldrich. Ammonium hydroxide (NH_4_OH) and 1N hydrochloric acid (HCl) were purchased from Fisher Scientific. Ethanol (EtOH) was purchased from Decon Labs, Inc. Phosphate-buffered saline (PBS, 10X) ultrapure was purchased from Thermo Fisher. These chemicals were used without further purification. Millipore Synergy UV-R Ultrapure (Type 1) water (DI H_2_O) of resistivity = 18.2 MΩ was used in all aqueous solutions. Aqua regia solution was used to clean the stir bars. Piranha solution or RCA1 and 1N HCl was used to clean the glass and silicon slides. The cleaned slides were stored in EtOH until further use. All syntheses were performed in 20 mL disposable plastic scintillation vials purchased from Fisher Scientific. No further purification was done to chemicals before use. Silica microparticles (0.6 μm) were purchased from Bangs Laboratories, Inc. In 1 mL portions, the microparticles were centrifuged (4000xg, 10 min, 4°C) and washed 3 times with EtOH. The final pellet was redispersed in fresh EtOH to prepare a 7w/v% particle solution for use. Commercial cell-line sEVs were purchased at ATCC (i.e., A549 exosomes (CCL-185-EXM), LNCap exosomes (CRL-1740-EXM), and hTERT MSC exosomes (SCRC-4000-EXM). Commercial gold and silver SERS substrates were purchased from Silmeco.

### Small extracellular vesicles (sEVs) isolation from clinical samples

sEVs were isolated from biofluids without the need for pre-purification steps. Each device, comprising eight chips, accommodates eight micropipettes for high throughput sEV extraction, allowing the extraction of 400 μL biofluid in approximately 20 minutes. The micropipettes were backfilled with 1X PBS buffer using a 33-gauge Hamilton syringe needle and positioned on the PMMA substrate. To initiate the process, the tip side of each micropipette was injected with 50 μL of biofluid, while the base side chamber received 1X PBS. Subsequently, sEVs from the biofluids were captured at the tip by applying a 10 V/cm direct current (DC) for 10 minutes, followed by release in 25 μL 1X PBS through the reversal of the applied voltage for another 10 minutes. To enhance efficiency, 1 mL biofluids underwent simultaneous processing using three iDEP devices, with parallel aliquots employed for purifying sEVs in 500 μL PBS. The isolated sEVs were stored at - 80°C for subsequent analysis.

**Table 1.** The clinical sEVs, also known as exosomes, used in this work.

| Subject/<br>Sample ID | Sample<br>Type | Gender/<br>Age at<br>collection | Race | Primary<br>Diagnosis | Histological<br>Subtype | Source |
| --- | --- | --- | --- | --- | --- | --- |
| URS400/<br>URS400ST1SE1 | Serum | Female/<br>43 yrs | Black or<br>African<br>American | Kidney<br>Cancer | Clear Cell<br>Papillary Renal<br>Cell Carcinoma | University of<br>Cincinnati<br>Biorepository |
| ISERS10 ML<br>37363-04/<br>ISERS10 ML<br>37363-04 | Serum | - | - | Healthy<br>Donor | Single Donor<br>Human serum<br>of the Clot | Innovative<br>Research |

The study was carried out in accordance with the principles of the Declaration of Helsinki under an approved protocol of the institutional review board of the University of Cincinnati (IRB# 2015-2364). Written informed consent was obtained and documented for all patients; samples were deidentified for patient confidentiality. Two of the urine samples were from the same patient. sEVs harvested from hTERT-immortalized mesenchymal stem cells and A549 NSCLC cell lines were purchased from ATCC Inc. For the rest of the paper, the kidney cancer sample will be denoted as a cancer patient and the healthy donor as a healthy control.

### Gold seeds synthesis

Gold citrate-capped nanoparticles (12 nm, 0.1 nM) were synthesized via a modified Turkevich method.^53^ In brief, 100 mL of deionized (DI) water and 200 µL of 0.5 M HAuCl_4_ were vigorously stirred and brought to a boil for 10 s. Afterwards, 15 mL of 1% Na_3_Cit was added to the boiling solution. The solution continued to boil for 15–30 min, while the total volume was maintained. Afterward, the resulting gold citrate-capped nanoparticle (AuNPs) solution was cooled to room temperature (r. t.), filtered through a 0.22 µm nitrocellulose membrane, and stored at 4°C until ready for use. Extinction profile and transmission electron microscopy (TEM) images can be found in Figures S1 and S2.

### Nanostars synthesis

The silver-coated gold nanostars were synthesized following a modified Vo-Dinh group synthesis.^54^ Briefly, a plastic scintillation vial of 10 mL of DI H_2_O was stirred (700 rpm, r. t.), then 10 µL 1 N HCl, followed by 493 µL of 5.08 mM HAuCl_4_ were added to the vial. After 5 seconds, 100 µL of 0.1 nM (12 nm) AuNPs were added. After 10 seconds, AgNO_3_ (50 uL) of various concentrations (2 or 3 mM; samples named S10, and S15 relating to the AgNO_3_ final concentration, 10 µM and 15 µM, respectively) quickly followed by 50 µL of 0.1 M AA. After this step, gold nanostars (AuNS) were synthesized and ready for immediate use. For silver-coated gold nanostars (AuNS-Ag), additional steps are required. After the addition of 0.1 M AA, the solution was allowed to stir for 30 s. Lastly, 0.1 M AgNO_3_ (20, 50, or 100 µL; samples named Ag2, Ag5, or Ag10 relating to the various final concentrations, 0.20, 0.50, and 1.0 mM, respectively) was added, immediately followed by 10 µL of NH_4_OH. The resulting NS solution continued to stir for 60-90 s to ensure the silver coating stabilized the gold nanostars. The final solution was transferred and incubated at room temperature for 2 h, idle.

The aged AuNS-Ag were centrifuged in ∼5.5 mL portions (1500 xg, 15 min, 4°C) and concentrated by 10 times. The 10X concentrated s AuNS-Ag were stored at 4°C until ready for use. The AuNS and AuNS-Ag nanoparticles were characterized via UV-vis spectroscopy and transmission electron microscopy (TEM) (Figure S1-S3).

### Substrate fabrication

To prepare solid-state SERS substrates, microscope glass slides were used as the base. First, the microscope slides were cut to 8 x 25 mm. First, the slides were cleaned in base, NH_4_OH/H_2_O_2_/H_2_O (1:1:5, RCA1), for 1 h with gently swirling every 15 min. Then RCA1 solution was replaced with 1 N HCl and soaked for another hour. Lastly, the slides were washed three times with ultrapure water (18 MΩ), once in fresh EtOH, and stored in fresh EtOH until use. The clean glass slides were removed and allowed to air dry. Immediately after drying, 10 µL of washed 7w/v% silica microparticle in EtOH solution was dispensed onto the surface. The cleaned slide was rotated to form a silica coating on the surface. Afterward, a layer of silver was deposited on top of the silica via the Denton Discovery 24 Sputtering System at a base pressure of 1e^7^ to 6e^-8^ Torr, deposition pressure 5.38 – 5.32 mTorr, and deposition speed of 10 Å/s. A Crystal sensor was used to measure the silver thickness. Three different silver film thicknesses were tested (i.e., 50, 125, and 200 nm). The metallic film over nanospheres (MeFON) was characterized via SEM and reflectance spectroscopy (Figures S4 and S5).

Silicon wafers were used as the base for preparing colloidal nanoparticles (i.e., AuNPs, S15, S10-Ag2, S10-Ag5, and S10-Ag10) SERS substrates. First, the silicon wafers were cut to 4 x 8 mm. The slides were cleaned with acid HCl/H_2_O_2_ (4:1, piranha) for 1 h, with gentle swirls every 15 min. Then, the slides were washed three times with ultrapure water (18 MΩ), and 3 times with fresh EtOH, and stored in fresh EtOH until use. To attach the colloidal nanoparticles to the silicon surface, we applied 2 different methods: (1) drop cast and (2) incubation. For drop cast, first, the cleaned, cut silicon wafers were incubated in fresh 10% APTES at 70°C for 1h. Afterward, the APTES functionalized silicon was rinsed in with EtOH, DI H_2_O, then EtOH. The washed substrate was dried at room temperature (10 min). Then, 50 µL of the 10X concentrated nanoparticles were dispensed on top of the APTES functionalized silicon. The nanoparticles were dried onto the surface overnight before use. For the incubation method, we followed a modified version of our previous report protocol.^55^ In brief, the cleaned glass was incubated in fresh 10% APTES under the same conditions as the previous method. Afterward, the APTES fictionalized silicon wafer was rinsed with EtOH, DI H_2_O, EtOH, and then H_2_O. The rinsed slide was placed in a clean microcentrifuge tube with 10X concentrated nanostars and incubated in the nanostars solution for 2 h. Lastly, the resulting nanoparticles silicon substrates were quickly rinsed with DI H_2_O, and dried at r. t., and ready for use. Colloidal nanoparticles SERS substrates were characterized via scanning electron microscopy (SEM) imaging (Figure S2 and S3). Commercial gold (Au) and silver (Ag) substrates were purchased and used to compare with the lab-made substrates (Figure S6).

### SERS measurements

The commercial cell-line sEVs and clinical sEVs’ concentrations and sizes were evaluated via nanoparticle tracking analysis (Figure S7). From this, 30 µL of 10^9^ particles/mL of sEVs in 1X PBS was dispensed on SERS substrate and allowed to dry overnight, r. t. The Raman and SERS spectra were obtained using a lab-built Raman microscope. Inverted dried samples were tested on a 50 mW 633 nm laser (PD-LD) at 20x objective in a Nikon Eclipse Ti Raman microscope with MaxLine, RazorEdge Dichroic, EdgeBasic; Semrock filters, and a Princeton Instruments Pixis100 CCD coupled with Isoplane160. Data were collected via LightField software under 10-second exposures, 25 acquisitions, and a minimum of 50 spectra per substrate in a 0.25π mm^2^ area at 5mW power. Commercial sEVs from ATCC were tested on 4 substrates per sEV type. Clinically isolated sEVs were tested on 6 substrates per sEV type. For the colloidal substrate, cell-line sEVs and clinically isolated sEVs were tested on 2 substrates per sEV type.

The SERS spectra obtained from the solid-state substrates range contained 359-2767 cm^−1^ with 1024 pixels. Next, the spectra were smoothed and background-subtracted using a Savitzky–Golay filter, with a 40-point window and first-order polynomial, on MATLAB 2021b, followed by normalized by dividing it for its integral (area under the curve of all Raman peaks) to avoid bias due to concentration differences. For the SERS spectra obtained by the colloidal-state substrates, the spectra range from 550-2767 cm^−1^ with 950 pixels. Smoothing, background removal, and normalization were performed the same way.

### Spectral classification using predictive and explainable machine learning

The characterization of cell-line commercial sEVs spectra using the solid-state SERS substrate was refined through a series of iterative tests, involving dimensionality reduction techniques and machine learning (ML) classifiers (python version 3.9.10). An Nvidia GeForce GTX 1060 GPU was used for analysis. The ultimate goal was to assess the performance of the final model in distinguishing between clinically isolated spectra of healthy and cancerous sEVs. Initially, Partial Least Squares Discriminant Analysis (PLS-DA) was conducted, determining the optimal number of PLS components through 10-fold cross-validation runs to maximize the accuracy (scikit package version 1.3). Applying a discriminant analysis, such as PLS-LDA, is also a common approach and was carried out using both linear (LDA) and quadratic (QDA) discriminant analysis. Additionally, Principal Component Analysis with Support Vector Machines (PCA/SVM) was employed to reduce the dimensionality of the dataset, which contained intensities of 984 pixels between 408-2720 cm^−1^, based on explained variance ratios. Finally, linear, cubic, and radial basis function kernels for SVM were applied to the reduced data set to compare classification performances.

In addition to the initial data transformation using PLS or PCA, the reduction of the high-dimensional dataset was also achieved through a process of eliminating peaks that were determined to be unnecessary for various trained classifiers (PyCaret package version 3.0.4). A comprehensive evaluation of 15 classifiers (Table S1) was carried out using a stratified 10-fold cross-validation approach. The scoring metrics collected include model accuracy, AUC, precision, recall, and F1-score. Taking the top 5 performing classifiers, based on the area under the curve (AUC), hyperparameter tuning was executed through 100 iterations, resulting in a total of 1,000 runs per classifier using RandomGridSearch, again basing optimization on the AUC. The best models after hyperparameterization were then selected to compute feature importance using SHapley Additive exPlanations (SHAP). Subsequently, the top 5% of pixels/features determined by each classifier were retained and employed for the best classifier(s) to identify the pixels that are most characteristic of changes between classes. Here, we randomly removed 30% of the observations as a test set and trained new models with the remaining 70% using the best parameterized algorithms. Final evaluation of top algorithms was taken from the scoring metrics of predicting the separated 30% of the data. The most successful model(s) were then applied to clinically isolated sEVs spectra to test the reliability of the prediction based on commercial cell-line samples for the clinical samples, and the interpretation of how the models predicted the test set was completed through SHAP beeswarm plots. This procedure was iteratively executed for the dataset utilizing solid-state and colloidal-state SERS substrates.

## Results and Discussion

Our long-term goal is to integrate SERS detection and the label-free microchip for sEVs purification in a single device. In this work, we optimize ML methods for SERS classification of sEVs. We performed the optimization on commercial cell-line sEVs and tested the methods on clinical sEVs samples extracted via the electrokinetic-based microchip. Both initial optimization and testing on clinical samples were performed using solid-state fabricated SERS substrates. These substrates (i.e., metallic film over nanospheres or MeFON) have superior SERS signal amplification and were used for initial optimization. However, these substrates are not easily integrated into a device. To this end, we then used the optimized ML methods to test the sEVs samples on the colloidal-state fabricated substrate (i.e., immobilized nanostars) demonstrating their efficacy. For both solid- and colloidal-state fabrication, we analyzed a series of substrate options to select the substrate with the highest signal, which was best suited for sEVs classification (Figure S8 and S9).

### SERS classification of cell-line derived sEVs

As test data for this analysis of classification methods, we tested different cell-line derived sEVs and controls. We collected SERS spectra on MeFON substrates (Figure 2A). To avoid concentration bias in the SERS spectra, the spectra were normalized by dividing them by their integral (area under the curve of all Raman peaks). In Figure S10, we show the individual spectra of sEVs compared to liposomes and PBS controls, which show observable differences. However, when comparing the specific sEVs spectra, there are only subtle variations (Figure 2B). The Raman Shift at 1000, 1030, 1200, and 1700 cm^-1^ are lower or higher normalized intensity for LNCap compared to A549 and hTERT. A549 and hTERT averaged spectra show slight differences at 1540 cm^-1^, where the signal is higher and more present in A549, and at 1320 cm^-1^, where a small signal can be observed in hTERT and not in A549. To classify these spectra, we tested several ML methods to extract the variance in these sEVs spectra and accurately classify the cell-line commercial sEVs. The goal of this study was to identify an ML method ideal for cancer diagnostics, defined as a method with high accuracy while conserving testable chemical insight. Importantly, while the spectra analyzed are from relatively homogenous samples of sEVs derived from a specific cell-line, there is significant variability associated with different SERS-substrate interaction with the sEVs, as well as with differences in sEVs composition. Figure S11 shows the single spectra for each sample (N > 200 spectra from at least 50 point-measurement on 4 substrates). This variability grants the use of complex multivariate analysis to classify these samples.

**Figure 2.**
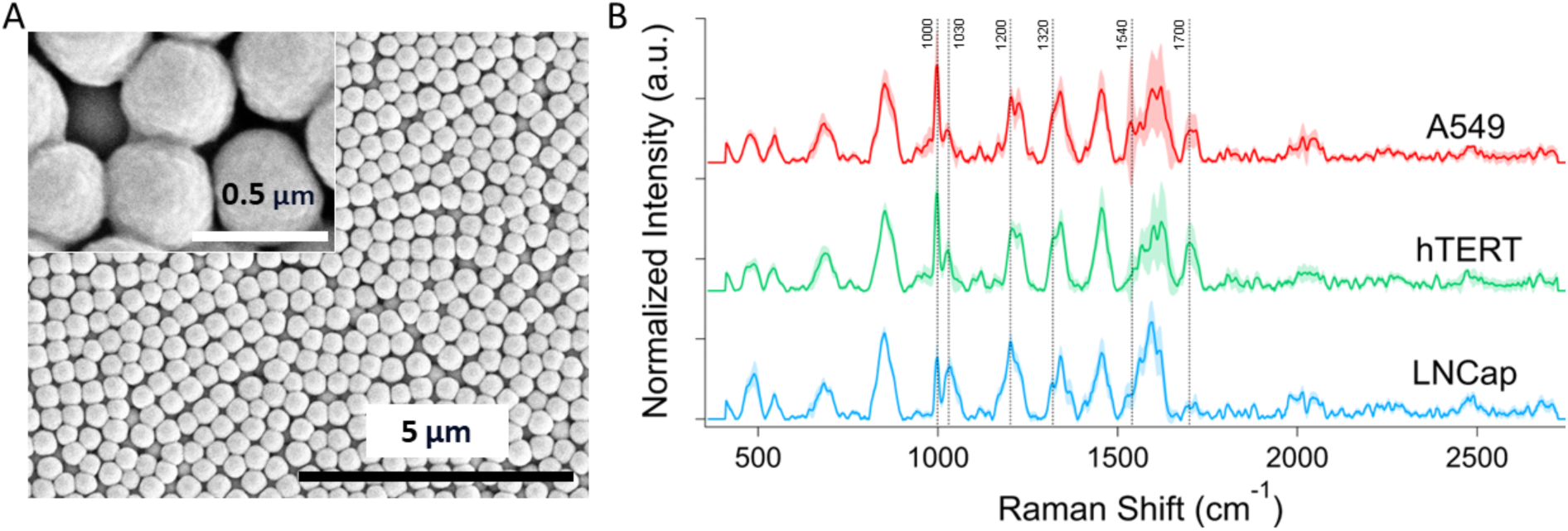
sEVs detection via label-free SERS on solid-state fabricated substrates. (a) SEM images of the SERS solid-state substrate metallic film over nanospheres (MeFON). (B) Averaged normalized SERS signals of commercial cell-line sEVs of A549 (red, top), hTERT (green, middle), and LNCap (blue, bottom).

For the initial benchmark classification, we use commonly adopted dimensionality reduction methods. We applied partial least squares discriminant analysis (PLS-DA) to the commercial cell-line sEVs spectral data, which included A549, hTERT, and LNCap samples measured using MeFON. The determination of the optimal number of components for reduction was based on maximizing the accuracy across 10 stratified cross-validation runs, testing a range from 1 to 40 components. The highest accuracy was achieved with 10 components (92.0%) (Figure S11). A visual examination of the first two components revealed a clear differentiation among the three classes, with a slight division observed with hTERT (Figure S11). We additionally attempted to apply linear (PLS-LDA) and quadratic (PLS-QDA) discriminant analysis with PLS to distinguish between the three classes. Here, we observed an increase in accuracy compared to PLS-DA, with only two components resulting in ∼94% average accuracy regardless of the discriminant analysis used (Figure S11). With 5 PLS components, the average accuracy for linear and quadratic discriminant analysis significantly increased to ∼99%. Notably, adding more than 5 components showed values near or exactly at 100% over all 10 k-folds, which demonstrates the high potential for overfitting when adding many PLS components. For this reason, our final analysis was completed using 5 PLS components, resulting in an average accuracy of 99.0% when applying LDA and 99.2% with QDA. The classification results from PLS-LDA and PLS-QDA were summarized as a confusion matrix (Figure 3 and S11). The models were able to predict each sEV accurately.

**Figure 3.**
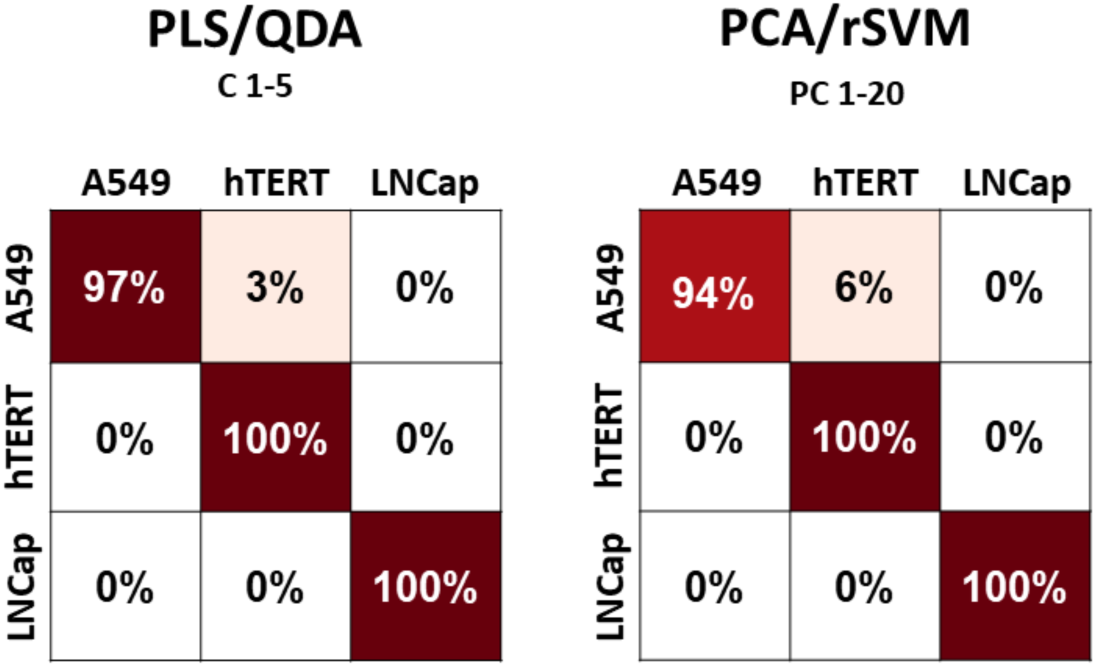
Confusion Matrix from PLS/QDA and PCA/ radial bias function (r) SVM classifications on commercial cell-line sEVs, A549, hTERT, and LNCap. PLS/QDA shows a higher classification among the sEVs compared to PLS/DA and PLS/LDA (Figure S11) with 3% misclassification of A549 as hTERT. PCA/SVM demonstrates 6% misclassification of A549 with hTERT, but overall high performance.

We employed a similar reduction method, principal component analysis (PCA), on the cell-line sEVs data to explore an alternative transformation of the original dataset. By retaining the first 20 components, we could explain less than 60% of the total variance present in the original dataset. Visualizing the first two components (PC1: 18.2%, PC2: 12.9% variance explained), we observed clustering of data points within each class, as depicted in (Figure S12). However, there was a small cluster with overlap between A549 and hTERT, which was expected given the similarity between the average spectra and their known protein expression of CD63 when compared to LNCap. To leverage as much variance as possible, we applied Support Vector Machines (SVM) with various kernels that enabled modeling in higher-dimensional spaces using the first 20 PC components. We found that the radial basis function kernel outperformed the polynomial and linear kernels, although all kernels produced scores above 90% (Table S3), and the confusion matrix summarized the classification results in Figure 3. There was a 6% misclassification for A549 sEVs and no misclassification for hTERT and LNCap sEVs from the validation set.

In summary, our analysis revealed PLS coupled with either LDA or QDA to be the most effective model for dimensionality reduction and classification of the commercial cell-line spectra data. PCA/SVM with the radial basis function kernel also showed successful application to the cell-line and clinical data. However, it’s worth mentioning that the PCA/SVM approach could only account for approximately 60% of the data’s total variance when using 20 components, which perpetuates the concerns about this method’s application on future data.

### Explainable ML algorithms comparison for sEVs classification

While PCA and PLS showed high accuracy in sEVs classification, the initial dimensionality reduction performed strongly reduces the interpretability of the classification model. To bolster our confidence in building a versatile model capable of classifying both the cell-line and clinical spectra, we transitioned from the initial dimensionality reduction approach to utilizing the normalized peak intensities in our classification (i.e., model applied to the Raman data with XAI). In this context, our objective was still to avoid the risk of overfitting a model with more features than observations. Additionally, our final goal is to produce a model for cancer diagnostics. Conserving interpretable Raman chemical information could be key in the clinical translation of this method, by offering a testable hypothesis for the classification model.

In this approach, we adopted a strategy of feature selection by identifying and eliminating features that were deemed nonessential for distinguishing healthy from cancer spectra. To determine the significance of each pixel or feature, we harnessed the power of SHAP, an XAI algorithm grounded in game theory that assigns weights to features based on their contributions to the model’s selection of a particular class. Through this process, we fit a finalized model that exclusively incorporated the most crucial features, those that had the greatest influence in discerning between the various spectra.

Our initial step involved conducting multiclass classification using a diverse set of 15 classifiers (Table S1). Here, we implemented a stratified k-fold cross-validation approach with a value of 10 and collected various metric scores. After rigorous evaluation, we identified the 5 top-performing classifiers, which included Light Gradient Boosting Machine (LGBM), Extra Trees (ET), CatBoost (CAT), Random Forest (RF), and Gradient Boosting (GB) based on their excellent average accuracy, AUC, recall, precision, and F1-score.

It is worth noting that each of these classifiers possesses a range of parameters whose optimal settings can heavily depend on the nature of the input data. To account for this variability, we conducted a thorough RandomGridSearch optimization process for the top 5 algorithms. This involved running RandomGridSearch for each algorithm 100 times, considering 1,000 runs per classifier, including cross-validation iterations. Table S4 provides comprehensive details on the final optimized settings for each algorithm.

Utilizing SHAP, we quantified the importance of every spectral pixel for each classifier. An additional facet that the SHAP analysis illuminates is how classifiers’ various learning architectures affect the quality of a specific dataset’s classification. Intriguingly, we uncovered a disparity in how bagging classifiers (ET and RF) and boosting classifiers (LGBM, CAT, and GB) were trained with the commercial cell-line data. The two bagging classifiers, which rely on averaging from multiple decision trees, recognized the top features as crucial for all three cell-culture sEVs. In contrast, the boosting classifiers, which learn through sequential predictions by implementing loss functions, predominantly identified features as important for only one or two sEVs. While it’s not surprising for features to play a pivotal role in characterizing one or two specific sEVs, observing that boosting algorithms were completely dependent on spectral pixels that are significant for only one or two sEVs led us to consider bagging algorithms as offering a more balanced classification approach (Figure 4). Given these insights, we decided to focus our final classification efforts on ET and RF classifiers. To further optimize and reduce dimensions, we employed SHAP to construct models using only the top 5% of the most important spectral pixels. Leveraging the top 5% of features (49 pixels), of which exact pixels differed slightly between ET and RF, we achieved remarkable results by using a 70/30 train and test split. We found RF reaching an accuracy of 96.2%, while ET demonstrated an exceptional accuracy of 99.7%, as detailed in Table S5 and Figure S13. We decided to move forward with ET.

**Figure 4.**
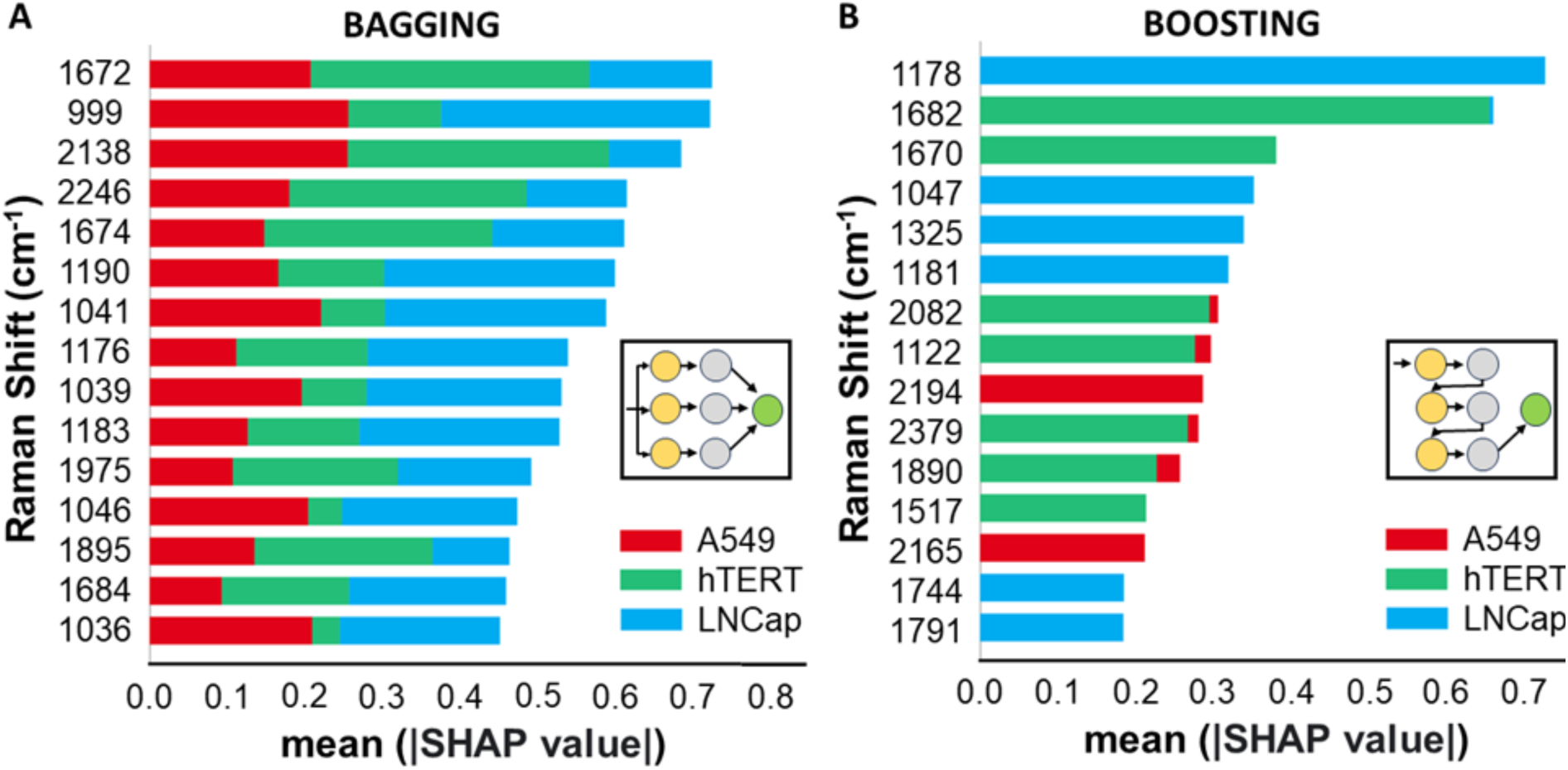
SHAP barplot from Bagging and Boosting Classifiers. SHAP barplot showing the most important features, calculated by the mean absolute SHAP value, and colored based of the feature’s importance per exosome for (A) the bagging algorithm Extra Trees (ET) Classifier and (B) the boosting algorithm Light Gradient Boosting (LGBM) Classifiers. Bagging algorithm demonstrates balanced contributions from each exosome in the classification. Boosting algorithm demonstrates unbalanced contribution from each exosome and leads to lower quality and bias towards one or two out of the three exosomes.

The ET confusion matrix for the classification of sEVs is shown in Figure 5, for direct comparison with PLS and PCA results. As can be observed, the accuracy of the test set is comparable with the dimensionality reduction methods, although slightly lower. However, the small decrease in accuracy is counterbalanced by a high degree of confidence in the model and by conserving the chemical insight through the classification process. As an example, the SHAP analysis shown in Figure 4 was used to determine the appropriate ML method (i.e., bagging) by looking at the model, rather than simply based on the accuracy of the results (effectively removing part of the black box). In addition, while PLS offers some visualization of the correlation between the classification model and the original data with the VIP scores (Figure S14), the actual PLS classification is not represented by the VIP scores, which reduces the interpretability of these results. With SHAP applied to the final separated test set, an XAI analysis, we can rank specific Raman bands based on their importance to characterize each sEVs, conserving a high degree of chemical insight, which is a major advantage of Raman.

**Figure 5.**
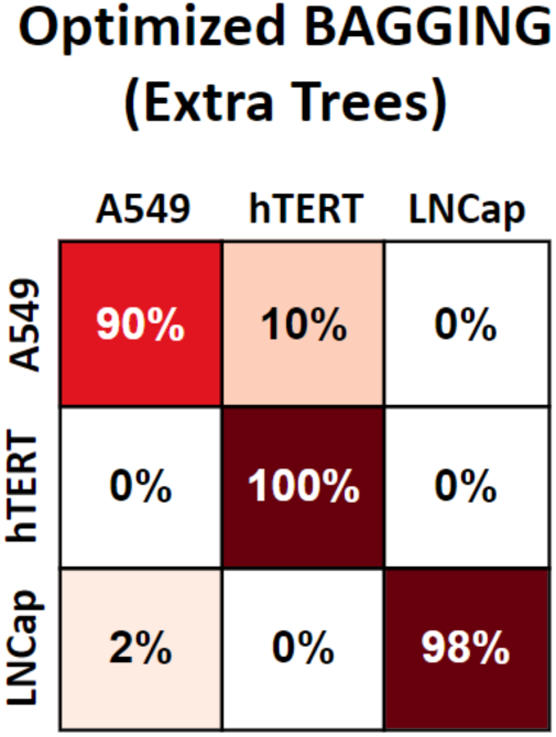
Confusion Matrix from the optimized Extra Trees Bagging algorithm classification on commercial cell-line sEVs of A549, hTERT, and LNCap. Compared to Figure 3, the Extra Trees demonstrate some misclassification compared to PLS/DA and PCA/SVM. However, Extra Trees can utilize SHAP, an XAI, to extrapolate and identify specific chemical insight to the classification.

### SERS classification of sEVs extracted from clinical samples

As a proof-of-concept for our target application, we collected SERS spectra from sEVs extracted from healthy and cancer blood samples (Figure 6A). Blood samples were collected in kidney-cancer and healthy individuals, and then sEVs were extracted via micropipette dielectrophoresis, which permitted to obtaining of clean sEVs samples.^33^ The sEVs were analyzed by the same process as the commercial cell-line sEVs, on MeFON SERS substrates by drying on the substrate surface before SERS measurements. To avoid concentration bias in the SERS spectra, the spectra were normalized by dividing them by their integral (area under the curve of all Raman peaks) (Figure 6B, S15). While these samples do not represent the variability of a pre-clinical study, each spectrum tested is unique and independent because of the multiple ways EVs can interact with the SERS substrate and because of the heterogenous EV composition of the samples. Figure S15 shows the single spectra for each sample (N > 300 spectra from at least 50 point-measurements on 6 substrates). Since we found consistent success with bagging algorithms in cell-line sEVs’ data classification, we exploited this analysis to evaluate their capability to distinguish between spectra obtained from clinical samples. We maintained the same parameters that were optimized for the cell-line dataset, ensuring a fair and valid comparison of the model’s performance across different datasets. The extended analysis of the clinical data exclusively used ET. For ET, a reduced features set comprising the top 5% (49 pixels) yielded an accuracy of 91.9% with the 70/30 data split. Figure 7 shows the confusion matrix results for the validation set and an area under the curve (AUC) of 0.95, with a sensitivity of 95.7% and a specificity of 97.9%. For completeness, we also show the results for the RF model (Figure S16), the second best-performing bagging model, including the PLS-DA/LDA/QDA (Figure S17, Table S6) and PCA/SVM (Figure S18, Table S7) models. Overall, we found that it was necessary to include more PLS components in order to reach the same level of accuracy as the cell-line sEVs samples. PLS-DA had an average accuracy of 92.1% using 11 PLS components, and both PLS-LDA and PLS-QDA both scored 99.7% on average also using 11 PLS components. As for PCA/SVM, the first 20 components explain less than 60% of the data’s total variance, which perpetuates the concerns about this method’s application on future data. Both PLS and PCA approaches are not highly statistical. Though ET’s classification accuracy is slightly lower, it is an explainable learning method. This slight difference can be counterbalanced by conserving the chemical insights and applicability of SHAP to identify and extrapolate the most important insights in the overall classification.

**Figure 6.**
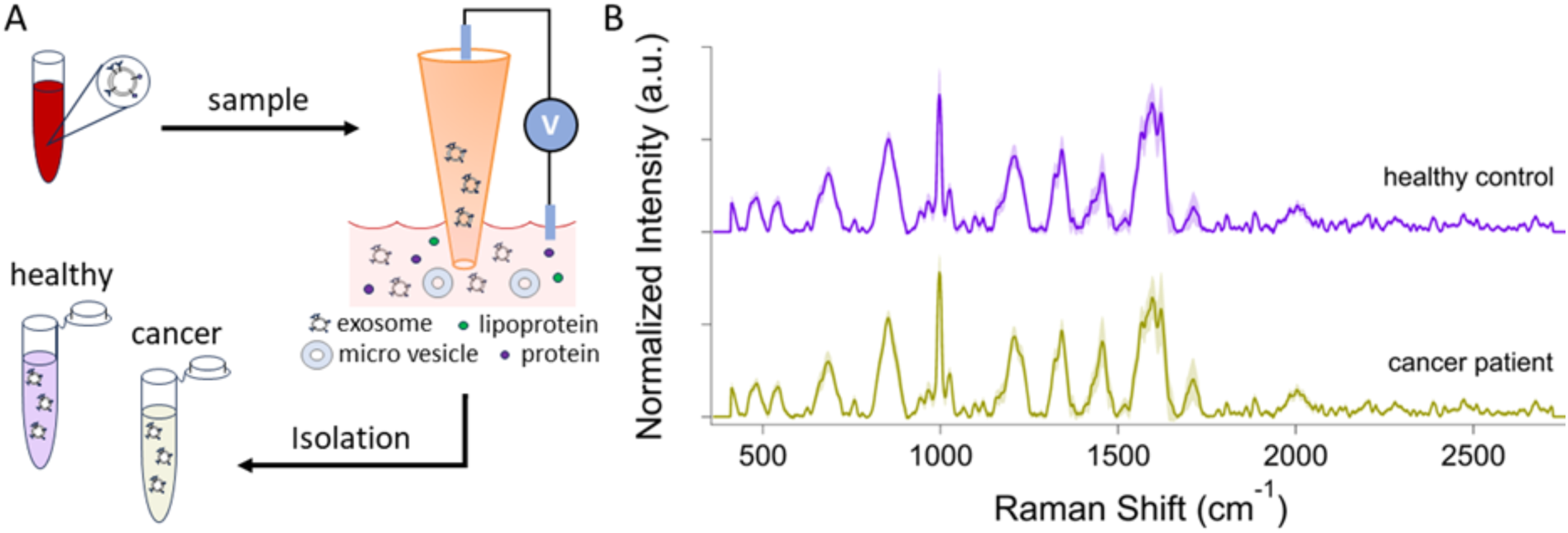
(A) Schematic of clinical sEVs-isolation via micropipette dielectrophoresis. (B) Averaged normalized SERS signals of clinical sEVs from healthy control (purple, top) and cancer patient (yellow, bottom).

**Figure 7.**
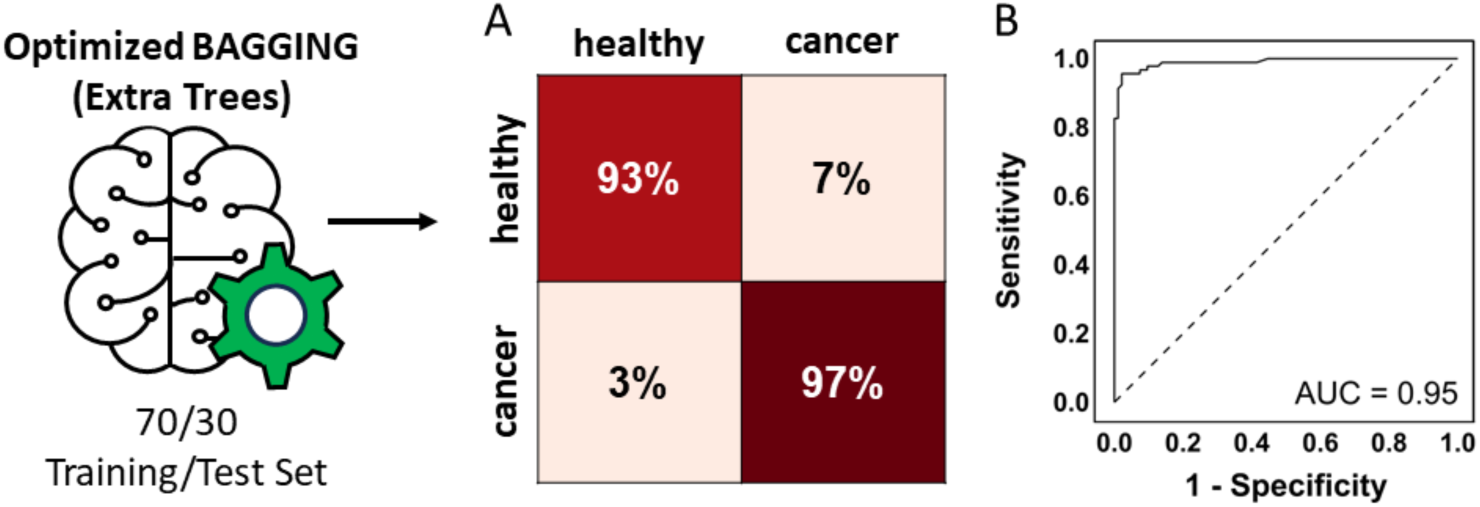
The application of the Optimized ML approach of Extra Trees Bagging on clinical sEVs, as a proof of concept, and the diagnostic accuracy. (A) The confusion matrix from the clinical sEVs, heathy control (healthy) and cancer patient (cancer) show high classification, and the (B) ROC Curve had an area under the curve (AUC) of 0.95.

### Important Raman spectral features from SHAP

From the previous section, the ET classifier was employed on the clinical sEVs. SHAP was further used to evaluate the ET’s performance by calculating the contributions of each spectral pixel to the model’s prediction of the final separated test set. This process allows specific identification of Raman signals that significantly influence the classification model. Figure 8A shows a SHAP beeswarm plot of Raman shift (Raman bands) and relative SHAP values. We limit the plot to the top 5% pixel based on SHAP importance. We also bracketed together pixels/shifts that are relative to the same peak in the spectrum for clarity. High absolute SHAP values signify more important Raman shifts for the classification among the top 5%. Figure 8A also shows the colors ranging from red to blue express whether the signals are on average higher or lower, respectively. Figure 7B highlights the 5% important Raman shift in the averaged spectra and with the most significant Raman shifts in red, to get a clear representation of the model on the collected spectra.

**Figure 8.**
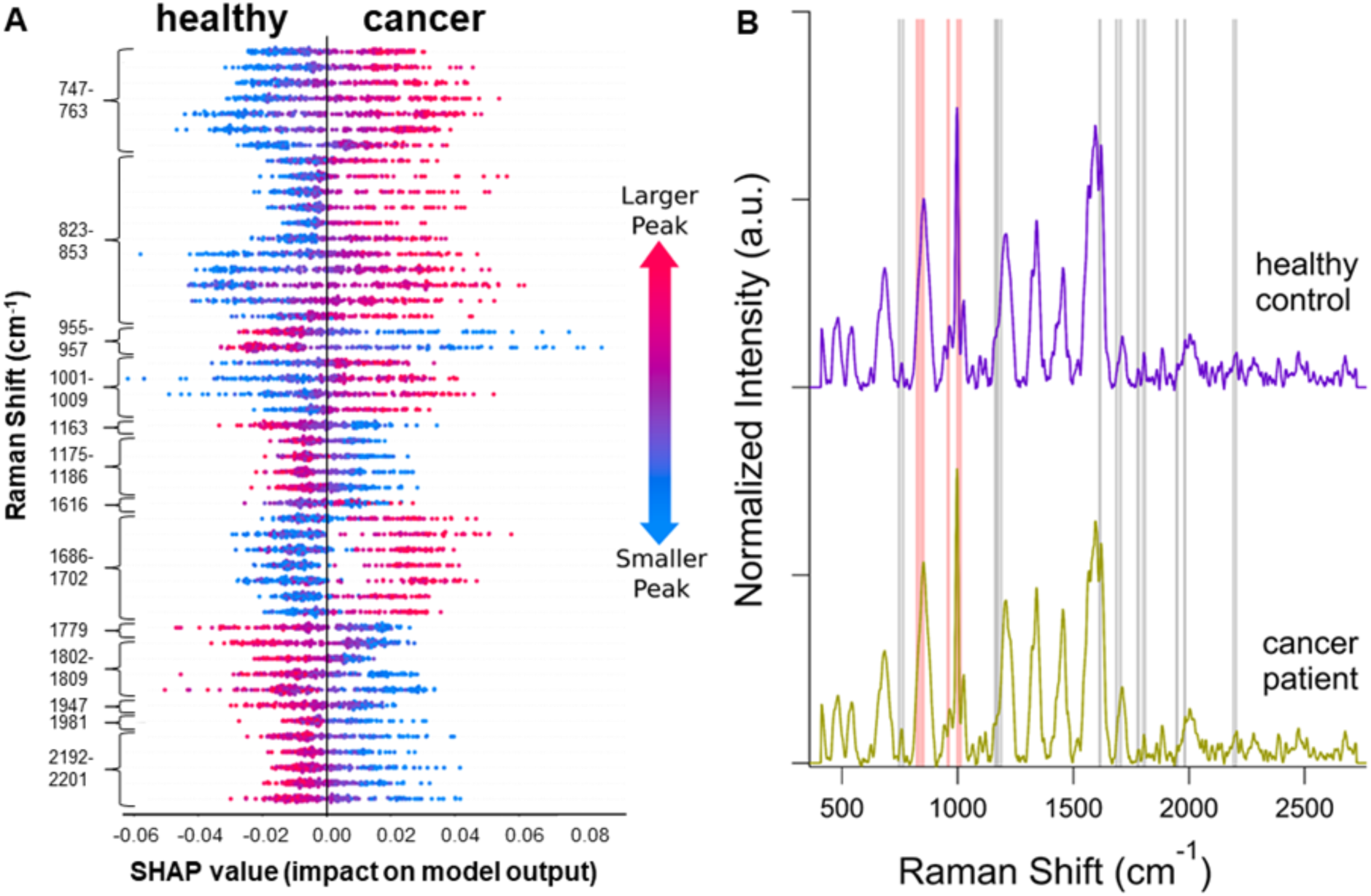
SHAP beeswarm plot (A) from the Optimized ML approach of Extra Trees Bagging on clinical sEVs and the results connection to the spectra (B). The SHAP beeswarm plot shows the top 5% of the most important Raman Shifts in the classification of the clinical sEVs on MeFON. (B) Averaged normalized SERS signals of clinical sEVs from healthy control (purple, top) and cancer patient (yellow, bottom). The gray and red highlighted regions are the top 5% most important in the classification. The red regions are the highest influence among the top 5%.

To better showcase the chemical information offered by SERS and SHAP analysis, we summarize the top 5% features in the ET classification in Table 2 which includes the band assignment and relative chemical component. The Raman features with the highest SHAP values (red regions in spectrum, Figure 7B) were attributed to ring system from Tyrosine (Tyr, 823-853 cm^-1^), C-C stretching (955-957 cm^-1^) likely from the α-helix backbone in proteins, and Phenylalanine (Phe, 1001-1009 cm^-1^). Tyr- and Phe-associated regions display a high-intensity Raman peak in the cancer spectra and lower intensity in the healthy spectra (as determined from the SHAP plot, Figure 7A). The region associated with C-C stretching shows the opposite behavior, with intensity higher in the healthy spectra. In addition, higher Raman intensity was from the Tryptophan’s ring system (Trp, 747-763 cm^-1^) and Amide I (1686-1702 cm^-1^) were observed for the cancer spectra. Other features highly significant for the model were assigned to Tyr ring breath (1163 cm^-1^), C-H bend (1175-1186 cm^-1^), and C=C stretching (1616 cm^-1^) from Tyr, Phe, and proteins. The Raman shift of other features may be assigned to C=O, C-H and C≡C stretching. These latter spectral regions had lower Raman intensities in the cancer spectra and higher intensities in the healthy spectra. Overall, our results suggest Raman signal from Tyr, Phe, Trp, and protein significantly contributed to the model’s prediction. In our results obtained from cell-line derived sEVs (Figure 4), we can see similar results with protein-associated features being the most important for model prediction (specifically, Phe and Amide I). This agreement determined using the SHAP analysis is a further validation of the robustness of this model.

**Table 2.** Important Raman features that made significant contributions to clinical sEVs classification via Extra Trees (ET) on metallic film over nanospheres (MeFON).

| Raman Shift (cm<sup>-1</sup>) | Assignment | Molecular components | Reference |
| --- | --- | --- | --- |
| 747-763 | symmetric ring breath | Trp, proteins | 56, 57 |
| 823-853 | v(C–C) proline, ring breath | Tyr, proteins | 56, 57 |
| 955-957 | v(C-C) stretching | α-Helix backbone, proteins | 56, 57 |
| 1001-1009 | v(C-C) skeletal, ring breath | Phe, proteins | 56, 57 |
| 1163 | ring breath | Tyr, proteins | 57 |
| 1175-1186 | v(C-H) bend | Tyr, Phe, proteins | 56, 57 |
| 1616 | v(C=C), stretching | Tyr, Trp, proteins | 56, 57 |
| 1686-1702 | Amide I, turn and bands | proteins | 56, 57 |
| 1779, 1802-1809 | v(C=O) | lipids | 58 |
| 1947, 1981 | v(C-H) | unassigned | - |
| 2192-2201 | v(C≡C) | lipids | 59 |

### sEVs classification on SERS substrates fabricated with colloidal nanostars

To extend our results toward our long-term goal, we used the optimized ML methods to examine SERS spectra obtained from substrates fabricated with colloidal nanoparticles. These substrates were fabricated by incubating functionalized silicon substrates in a suspension of concentrated silver-coated gold nanostars (NS) (Figure 9A), previously optimized (Figure S9). This fabrication method is more suitable to be added to a potential sEVs-extraction device to automate this type of analysis. However, colloidal fabrication of SERS substrates is often characterized by lower signal enhancements, which could reduce classification accuracy. To determine feasibility, we tested the classification of clinical samples with these substrates. We applied the ET classifier on the normalized spectra to find similar scoring metrics found with the solid-state substrate classification. Results for the classification are shown in Figure 9.

**Figure 9.**
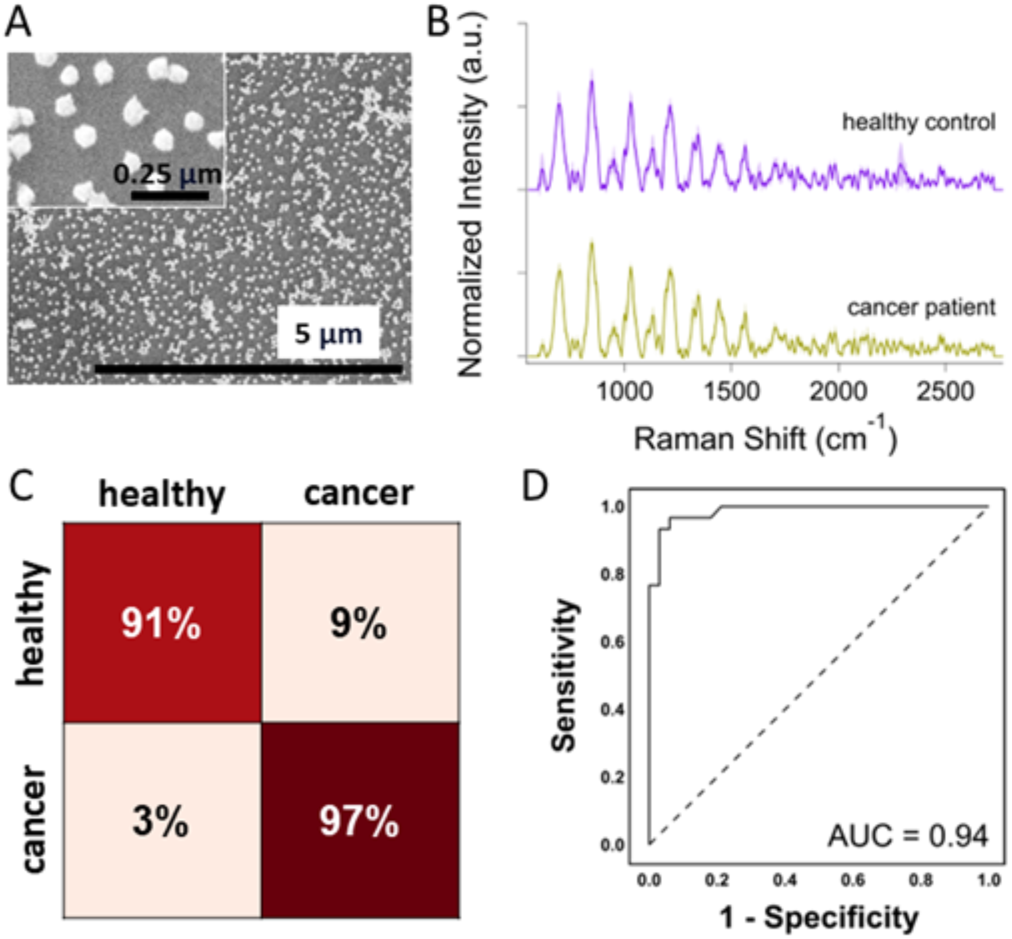
sEVs detection via label-free SERS on colloidal-state fabricated substrates and diagnostic accuracy. (A) SEM images of silver-coated gold nanostars (NS) on silicon. (B) Averaged normalized SERS signals of clinically isolated sEVs from human blood. The Extra Trees (ET) Bagging classification performance is summarized in the confusion matrix (C) and ROC curve (D).

Figure 9B shows the spectra obtained with these substrates on clinical sEVs samples. These spectra were significantly different from previous results with MeFON, mostly due to the different background signal and signal-to-background ratio for the sEVs signal. A blank spectrum is shown in Figure S19), to compare with the sEVs signal from the figure. Figure 9C, D, the ET classification shows high accuracy for the identification of sEVs from cancer patients, as shown in the confusion matrix of the validation set and in the ROC curve, showing an AUC of 0.94. The accuracy for NS substrates was below what was observed using MeFON. In addition, compared to results obtained with MeFON substrates, the SHAP analysis showed an overall weaker classification, with the top 5% exhibiting smaller magnitudes of importance overall (Figure S20). More importantly, the top 5% spectral peaks collected differed in regions as well as trends found in the MeFON, solid-state substrate (Figure 8). This result is likely due to the lower intensity of peaks associated with the sEVs, with some peaks not observed with NS substrates. Overall, these results sEVs the superiority of MeFON substrates but also demonstrate the feasibility of performing classification with NS substrates. We believe that further optimization of NS substrates could lead to robust SERS classification in sEVs-extraction devices.

## Conclusion

In summary, we study the use of ML methods for sEVs, exosomes, classification from sEVs derived from cell-lines and extracted from clinical samples. This study was performed by first evaluating different classification methods (PLS-DA/LDA/QDA, PCA/radial SVM, and ET/SHAP) for cell-line-derived sEVs. These studies show high classification accuracy for all methods but revealed how PLS and PCA are not explainable methods, thus making it difficult to bring the gap between of ML models to human comprehension and trust in clinical settings. The results show the superiority of XAI, which through SHAP analysis permitted to selection of the most robust model (bagging classifer, ET) and to directly relate spectral features to the classification model, avoiding overfitting and bias. As a proof of concept, the optimized ET/SHAP model approach was further used on sEVs extracted from clinical samples, showing high diagnostic accuracy. These results show the potential for SERS analysis to be used for non-invasive cancer diagnostics. Finally, we tested clinical samples on substrates fabricated with NS, proving that the classification was possible also on this type of cheap and versatile substrates. In conclusion, we demonstrated the integration of label-free microchip with SERS, including with colloidal NS that can easily be integrated in a potential device. We also showcased the use of XAI to perform diagnostic analysis on complex SERS data of clinical samples, demonstrating how this approach enables optimization and interpretation of the model due to its direct association to the input spectral data.

## Acknowledgments

This research was supported by the University of Cincinnati’s University Research Council (URC) Graduate Student Stipend and Research Cost Program for Faculty— Student Collaboration: Physical Sciences and Engineering. The authors acknowledge funding by the National Science Foundation NSF CAREER ECCS (2046037), CincyTech, LLC/Ohio Development Services Agency (ODSA)/Entrepreneurial Services Program (ESP) (TECG2020041). The human biospecimens and associated clinical data were provided by the University of Cincinnati Cancer Center Biospecimen Shared Resource. In addition, the authors thank Ronald G. Flenniken of Engineering at the University of Cincinnati for depositing the metallic silver film on the nanosphere substrates used in the study. In addition, we thank Jessica Webster of the Pathology Research at Cincinnati Children’s, Mahnoosh Khosravifar, and Melodie A. Fickenscher of CEAS at the University of Cincinnati for their knowledge and advice on TEM or SEM.

## References

1. Siegel, R. L.; Miller, K. D.; Fuchs, H. E.; Jemal, A. “Cancer statistics 2022”. Ca Cancer J Clin 2022, 71 (1), 7–33.

2. Munir, K.; Elahi, H.; Ayub, A.; Frezza, F.; Rizzi, A. “Cancer diagnosis using deep learning: a bibliographic review.” Cancers 2019, 11 (9), 1235.

3. Wilkinson, A. N. “Cancer diagnosis in primary care: Six steps to reducing the diagnostic interval.” Can Fam Physician. 2021, 67 (4), 265–268.

4. Atlihan-Gundogdu, E.; Ilem-Ozdemir, D.; Ekinci, M.; Ozgenc, E.; Demir, E. S.; Sánchez-Dengra, B.; González-Alvárez, I. “Recent developments in cancer therapy and diagnosis.” J. Pharm. Investig. 2020, 50, 349–361.

5. Litwin, M. S., Tan, H.-J. “The diagnosis and treatment of prostate cancer: a review.” JAMA 2017, 317 (24), 2532–2542.

6. Issa, I. A.; Noureddine, M. “Colorectal cancer screening: An updated review of the available options.” World J. Gastroenterol 2017, 23 (28), 5086–5096.

7. Rijavec, E.; Coco, S.; Genova, C.; Rossi, G.; Longo, L.; Grossi, F. “Liquid biopsy in non-small cell lung cancer: highlights and challenges.” Cancers 2019, 12 (1), 17.

8. Cheng, F.; Su, L.; Qian, C. “Circulating tumor DNA: a promising biomarker in the liquid biopsy of cancer.” Oncotarget 2016, 7 (30), 48832.

9. Di Capua, D.; Bracken-Clarke, d.; Ronan, K.; Baird, A.-M.; Finn, S. “The liquid biopsy for lung cancer: state of the art, limitations and future developments.” Cancers 2021, 13 (16), 3923.

10. Ansari, J.; Yun, J. W.; Kompelli, A. R.; Moufarrej, Y. E.; Alexander, J. S.; Herrera, G. A.; Shackelford, R. E. “The liquid biopsy in lung cancer.” Genes & cancer 2016, 7 (11-12), 355.

11. Thery, C. “Exosomes: secreted vesicles and intercellular communications” F1000 Bio. Rep. 2011, 3, 15.

12. Yu, D.; Li, Y.; Wang, M.; Gu, J.; Xu, W.; Cai, H.; Fang, X.; Zhang, X. “Exosomes as a new frontier of cancer liquid biopsy.” Molecular Cancer 2011, 21 (1), 1–33.

13. Thakur, A.; Parra, D. C.; Motallebnejad, P.; Brocchi, M.; Chen, H. J. “Exosomes: Small vesicles with big roles in cancer, vaccine development, and therapeutics.” Bioactive materials 2022, 10, 281–294.

14. Li, J.; Li, Y.; Li, P.; Zhang, Y.; Du, L.; Wang, Y.; Zhang, C.; Wang, C. “Exosome detection via surface-enhanced Raman spectroscopy for cancer diagnosis.” Acta Biomaterialia 2022, 144, 1–14.

15. Sheth, M.; Esfandiari, L. “Bioelectric dysregulation in cancer initiation, promotion, and progression.” Frontiers in Oncology 2022, 12, 846917.

16. Doyle, L. M.; Wang, M. Z. “Overview of Extracellular Vesicles, Their Origin, Composition, Purpose, and Methods for Exosome Isolation and Analysis” Cells 2019, 8 (7), 727.

17. Rana, A.; Zhang, Y.; Esfandiari, L. “Advancements in microfluidic technologies for isolation and early detection of circulating cancer-related biomarkers.” Analyst 2018, 143 (13), 2971–2991.

18. Shi, L.; Esfandiari, L. “Emerging on-chip electrokinetic based technologies for purification of circulating cancer biomarkers towards liquid biopsy: A review.” Electrophoresis 2022, 43 (1-2), 288–308.

19. Lewis, J.M.; Vyas, A. D.; Qiu, Y.; Messer, K. S.; White, R.; Heller, M. J. “Integrated Analysis of Exosomal Protein Biomarkers on Alternating Current Electrokinetic Chips Enables Rapid Detection of Pancreatic Cancer in Patient Blood.” ACS Nano 2018, 12 (4), 3311–3320.

20. Chen, Y.S.; Ma, Y.-D.; Chen, C.; Shiesh, S.-C.; Lee, G.-B. “An integrated microfluidic system for on-chip enrichment and quantification of circulating extracellular vesicles from whole blood.” Lab Chip 2019, 19 (19), 3305–3315.

21. Ayala-Mar, S.; Perez-Gonzalez, V. H.; Mata-Gómez, M. A.; Gallo-Villanueva, R. C.; González-Valdez, J. Anal. Chem. 2019, 91 (23), 14975–14982

22. Pethig, R. “Review article-dielectrophoresis: status of the theory, technology, and applications.” Biomicrofluidics 2010, 4 (2), 022811.

23. Ozuna-Chacon, S.; Lapizco-Encinas, B. H.; Rito-Palomares, M.; Martínez-Chapa, S. O.; Reyes-Betanzo, C. “Performance characterization of an insulator-based dielectrophoretic microdevice.” Electrophoresis 2008, 29 (15), 3115–3122.

24. Konoshenko, M.Y.; Lekchnov, E. A.; Vlassov, A. V.; Laktionov, P. P. “Isolation of Extracellular Vesicles: General Methodologies and Latest Trends.” Biomed Res Int 2018, 2018, 8545347.

25. Shi, L.; Rana, A.; Esfandiari, L. “A low voltage nanopipette dielectrophoretic device for rapid entrapment of nanoparticles and exosomes extracted from the plasma of healthy donors.” Sci Rep 2018, 8 (1), 6751.

26. Shi, L.; Kuhnell, D.; Borra, V. J.; Langevin, S.M.; Nakamura, T., Esfandiari, L. “Rapid and label-free isolation of small extracellular vesicles from biofluids utilizing a novel insulator based dielectrophoretic device.” Lab Chip. 2019, 19 (21), 3726–3734.

27. Sharma, M.; Sheth, M.; Poling, H. M.; Kuhnell, D.; Langevin, S. M.; Esfandiari, L. “Rapid purification and multiparametric characterization of circulating small extracellular vesicles utilizing a label-free lab-on-a-chip device*.”* Sci Rep 2023, 13, 18293.

28. Shi, L.; Esfandiari, L. “A label-free and low-power microelectronic impedance spectroscopy for characterization of exosomes.” Plos one 2022, 17 (7), e0270844.

29. Shi, L.; Esfandiari, L. “An electrokinetically-driven microchip for rapid entrapment and detection of nanovesicles. Micromachines 2020, 12 (1), 11.

30. Zhang, Y.; Murakami, K.; Borra, V. J.; Ozen, M. O.; Demirci, U.; Nakamura, T.; Esfandiari, L. “A label-free electrical impedance spectroscopy for detection of clusters of extracellular vesicles based on their unique dielectric properties.” Biosensors 2022, 12 (2), 104.

31. Špilak, A.; Brachner, A.; Kegler, U.; Neuhaus, W.; Noehammer, C. “Implications and pitfalls for cancer diagnostics exploiting extracellular vesicles.” Advanced Drug Delivery Reviews 2021, 175, 113819.

32. Shin, H.; Jeong, H.; Park, J.; Hong, S.; Choi, Y. “Correlation between Cancerous Exosomes and Protein Markers Based on Surface-Enhanced Raman Spectroscopy (SERS) and Principal Component Analysis (PCA).” ACS Sensors 2018, 3 (12), 2637–2643.

33. Yan, Z.; Dutta, S.; Liu, Z.; Yu, X.; Mesgarzadeh, N.; Ji, F.; Bitan, G.; Xie, Y.-H. “A Label-Free Platform for Identification of Exosomes from Different Sources.” ACS Sensors 2019, 4 (2), 488–497.

34. Rojalin, T.; Koster, H. J.; Liu, J.; Mizenko, R. R.; Tran, Di.; Wachsmann-Hogiu, S.; Carney, R. P. ACS Sensors 2020, 5 (9), 2820–2833.

35. Xiong, H.; Huang, Z.; Yang, Z.; Lin, Q.; Yang, B.; Fang, X.; Liu, B.; Chen, H.; Kong, J. “Recent progress in detection and profiling of cancer cell-derived exosomes.” Small 2021, 17, 2007971.

36. Guerrini, L.; Garcia-Rico, E.; O’Loghlen, A.; Giannini, V.; Alvarez-Puebla, R. A. Surface-Enhanced Raman Scattering (SERS) Spectroscopy for Sensing and Characterization of Exosomes in Cancer Diagnosis. Cancers 2021, 13, 2179.

37. Koster, H. J.; Rojalin, T.; Powell, A.; Pham, D.; Mizenko, R. R.; Birkeland, A. C.; Carney R. P. Nanoscale 2021, 13, 14760–14776.

38. Xie, Y.; Su, X.; Wen, Y.; Zheng, C.; Li, M. “Early-Stage Lung Cancer Diagnosis by Deep Learning-Based Spectroscopic Analysis of Circulating Exosomes.” Nano Letters 2022, 22 (19), 7910–7918.

39. Kazemzadeh, M.; Martinez-Calderon, M.; Paek, S. Y.; Lowe, M.; Aguergaray, C.; Xu, W.; Chamley, L. W.; Broderick, N. G. R.; Hisey, C. L. ACS Sensors 2022, 7 (6), 1698–1711.

40. Diao, X.; Li, X.; Hou, S.; Li, H.; Qi, G.; Jin, Y. “Machine Learning-Based Label-Free SERS Profiling of Exosomes for Accurate Fuzzy Diagnosis of Cancer and Dynamic Monitoring of Drug Therapeutic Processes.” Analytical Chemistry 2023, 95 (19), 7552–7559.

41. Wang, J.; Wuethrich, A.; Sina, A.A.I.; Lane, R.E.; Lin, L.L.; Wang, Y.; Cebon, J.; Behren, A.; Trau, M. “Tracking extracellular vesicle phenotypic changes enables treatment monitoring in melanoma.” Science advances 2020, 6 (9), p.eaax3223.

42. Blake, N.; Gaifulina, R.; Griffin, L.D.; Bell, I.M.; Thomas, G.M.H. “Machine Learning of Raman Spectroscopy Data for Classifying Cancers: A Review of the Recent Literature.” Diagnostics 2022, 12, 1491.

43. Kanno, N.; Kato, S.; Ohkuma, M.; Matsui, M.; Iwasaki, W.; Shigeto, S. “Machine learning-assisted single-cell Raman fingerprinting for in situ and nondestructive classification of prokaryotes.” iScience 2021, 24 (9), 102975.

44. Pavillon, N; Smith, N. “Deriving Accurate Molecular Indicators of Protein Synthesis through Raman-Based Sparse Classification.” Analyst 2021, 146, 3633.

45. Kothari, R.; Jones, V.; Mena, D.; Reyes, V. B.; Shon, Y.; Smith, J. P.; Schmozle, D.; Cha, P. D.; Lai, L.; Fong, Y., Storrie-Lombardi, M. C. “Raman spectroscopy and artificial intelligence to predict the Bayesian probability of breast cancer.” Sci Rep 2021, 11, 6482.

46. Shin, H.; Choi, B.H.; Shim, O.; Kim, J.; Park, Y.; Cho, S. K.; Kim, H. K.; Choi, Y. “Single test-based diagnosis of multiple cancer types using Exosome-SERS-AI for early stage cancers.” Nat Commun. 2023, 14, 1644.

47. Kim, S.; Choi, B. H.; Shin, H.; Kwon, K.; Lee, S. Y.; Yoon, H. B.; Kim, H. K.; Choi. Y. “Plasma Exosome Analysis for Protein Mutation Identification Using a Combination of Raman Spectroscopy and Deep Learning.” ACS Sensors 2023, 8 (6), 2391–2400.

48. Parlatan, U.; Ozen, M. O.; Kecoglu, I.; Koyuncu, B.; Torun, H.; Khalafkhany, D.; Loc, I.; Ogut, M. G.; Inci, F.; Akin, D.; Solaroglu, I.; Ozoren, N.; Unlu, M. B.; Demirci, U. “Label-Free Identification of Exosomes using Raman Spectroscopy and Machine Learning.” Small 2023, 19, 2205519.

49. Lussier, F.; Thibault, V.; Charron, B.; Wallace, G. Q.; Masson, J.-F. “Deep Learning and Artificial Intelligence Methods for Raman and Surface-Enhanced Raman Scattering.” TrAC 2020, 124, 115796.

50. Lundberg, S.M.; Erion, G.; Chen, H.; DeGrave, A.; Prutkin, J. M.; Nair, B.; Katz, R.; Himmelfarb, J.; Bansal, N.; Lee, S.-I. “From local explanations to global understanding with explainable AI for trees.” Nat Mach Intell 2020, 2, 56–67.

51. Hassija, V.; Chamola, V.; Mahapatra, A.; Singal, A.; Goel, D.; Huang, K.; Scardapane, S.; Spinelli, I.; Madmud, M.; Hussain, A. *“* Interpreting Black-Box Models: A Review on Explainable Artificial Intelligence.” Cogn Comput 2024, 16, 45–74.

52. Alabi, R.O.; Elmusrati, M.; Leivo, I.; Almangush, A.; Mäkitie, A. A. “Machine learning explainability in nasopharyngeal cancer survival using LIME and SHAP.” Sci Rep 2023, 13, 8984.

53. Turkevich, J.; Stevenson, P.C.; Hillier, J. “A Study of the Nucleation and Growth Processes in the Synthesis of Colloidal Gold.” Discuss. Faraday Soc. 1951, 11, 55–75.

54. Fales, A. M.; Yuan, H.; Vo-Dinh, T. “Development of Hybrid Silver-Coated Gold Nanostars for Nonaggregated Surface-Enhanced Raman Scattering.” J. Phys. Chem. C 2014, 118, 7, 3708–3715.

55. Vang, D.; Strobbia, P. “Analysis of nanostar reshaping kinetics for optimal substrate fabrication.” Applied Spectroscopy 2023. 77 (3), 270–280.

56. Kruglik, S. G.; Royo, F.; Guigner, J. M.; Palomo, L.; Seksek, O.; Turpin, P. Y.; Tatischeff, I.; Falcón-Pérez, J.M. “Raman tweezers microspectroscopy of circa 100 nm extracellular vesicles.” Nanoscale 2019, 11 (4), 1661–1679.

57. Movasaghi, Z.; Rehman, S.; Rehman, I. U. “Raman spectroscopy of biological tissues.” Applied Spectroscopy Reviews 2007, 42 (5), 493–541.

58. Kluz-Barłowska, M.; Kluz, T.; Paja, W.; Sarzyński, J.; Łączyńska-Madera, M.; Odrzywolski, A., Król, P.; Cebulski, J.; Depciuch, J. “FT-Raman data analyzed by multivariate and machine learning as a new methods for detection spectroscopy marker of platinum-resistant women suffering from ovarian cancer.” Scientific Reports 2023, 13 (1), 20772.

59. Taylor, R. W.; Benz, F.; Sigle, D. O.; Bowman, R. W.; Bao, P.; Roth, J. S.; Heath, G.R.; Evans, S.D.; Baumberg, J. J. “Watching individual molecules flex within lipid membranes using SERS.” Scientific Reports 2014, 4 (1), 5940.

