## supporting information for "Machine Learning Approaches in Label-Free Small Extracellular Vesicles Analysis with Surface-Enhanced Raman Scattering (SERS) for Cancer Diagnostics"

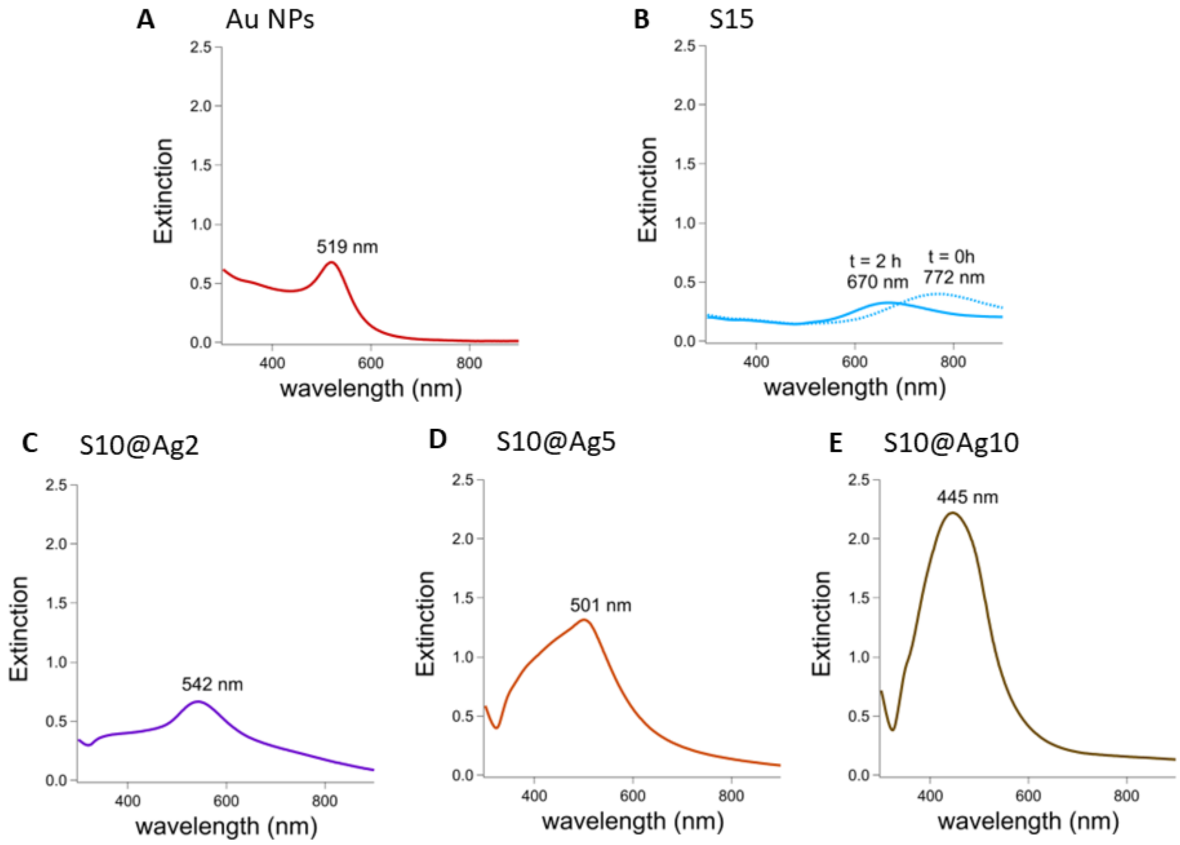


**Figure S1**. Ultraviolet-visible extinction spectra of (A) Au NPs, (B) S15 after synthesis and 2h, (C) S10@Ag2, (D) S10@Ag5, and (E) S10@Ag10.

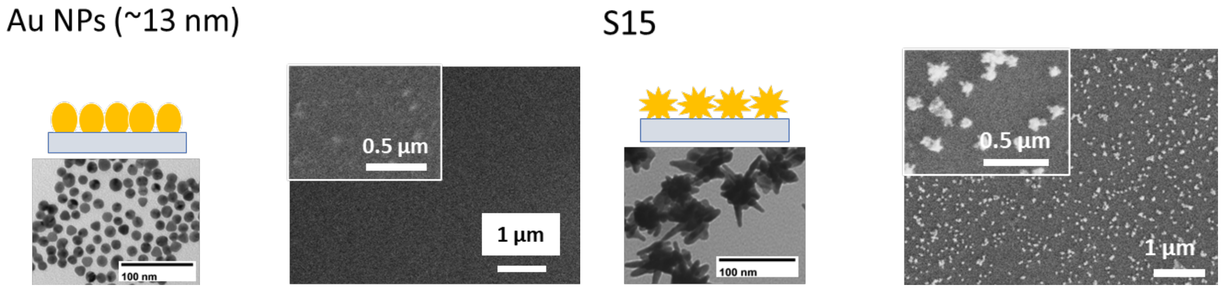


**Figure S2**. TEM images of colloidal nanoparticles of (left) Au NPs and (right) S15 and their SEM imaging of the nanoparticle on silicon.

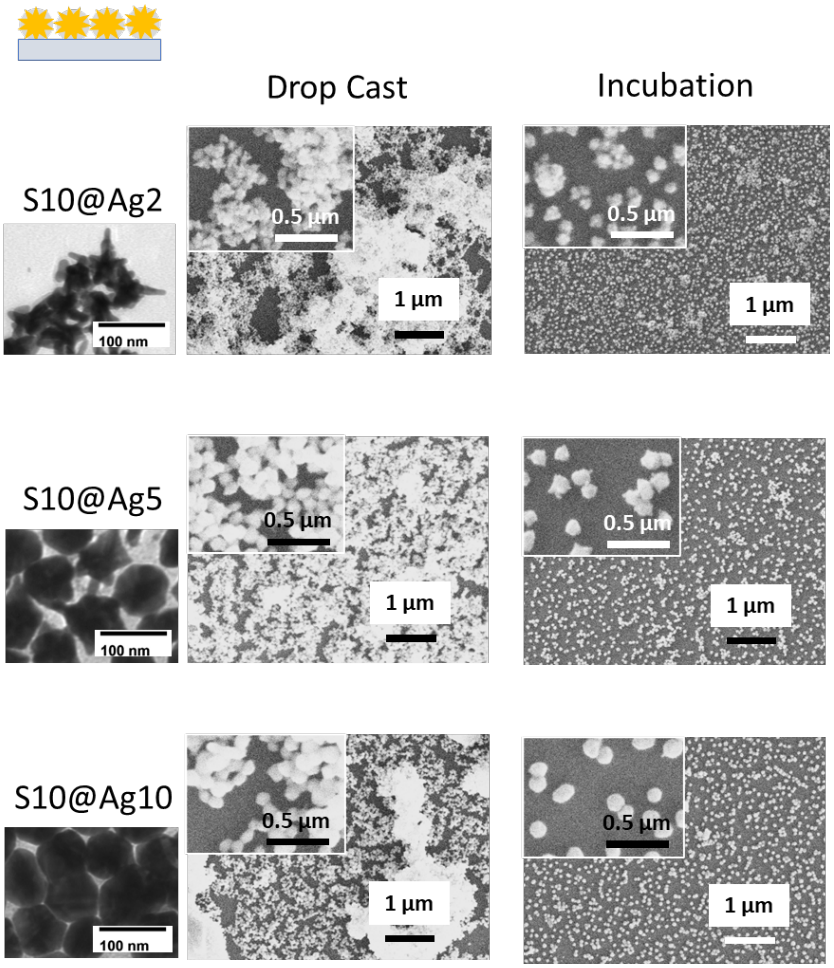


**Figure S3**. TEM imaging of colloidal S10@Ag2, S10@Ag5, S10@Ag10 nanoparticles and SEM imaging of the nanoparticle on silicon via drop cast and incubation.

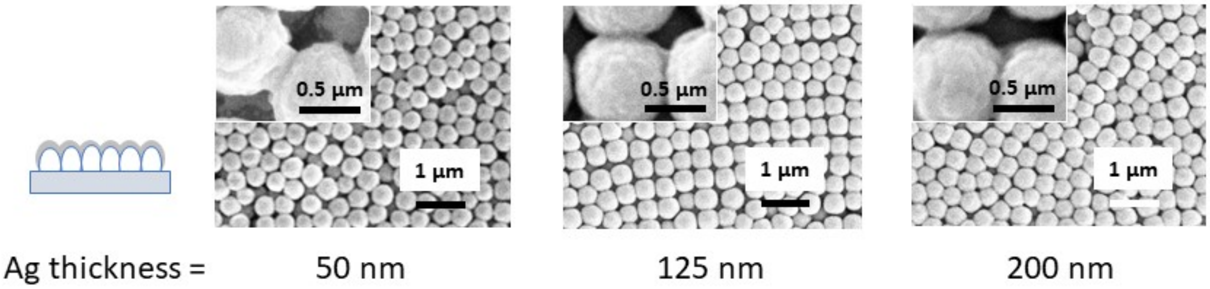


**Figure S4**. SEM images of Metallic Silver Film Over Silica Nanospheres (MeFON), solid-state fabricated, SERS substrates with 50, 125, and 200 nm Ag films.

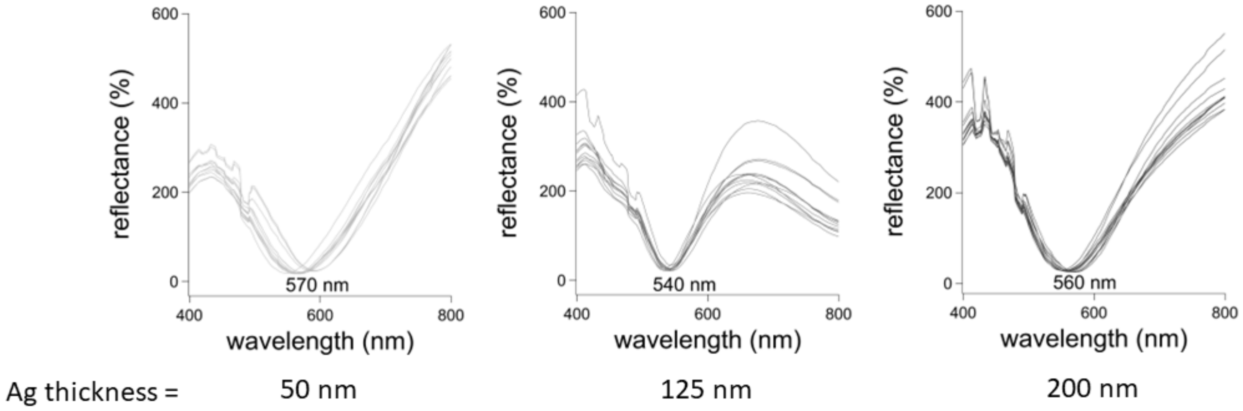


**Figure S5**. Reflectance spectra of Metallic Silver Film Over Silica Nanospheres (MeFON), solid-state fabricated, SERS substrates with 50, 125, and 200 nm Ag films.

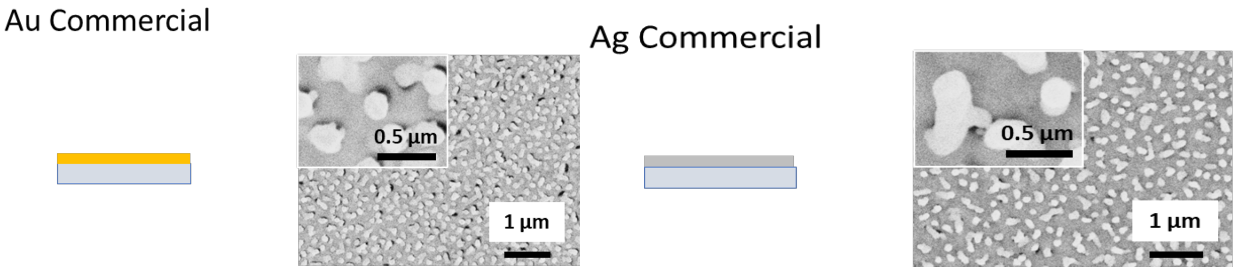


**Figure S6**. SEM images of the commercially available SERS substrate after 15 months of gold (Au, left) and silver (Ag, right).

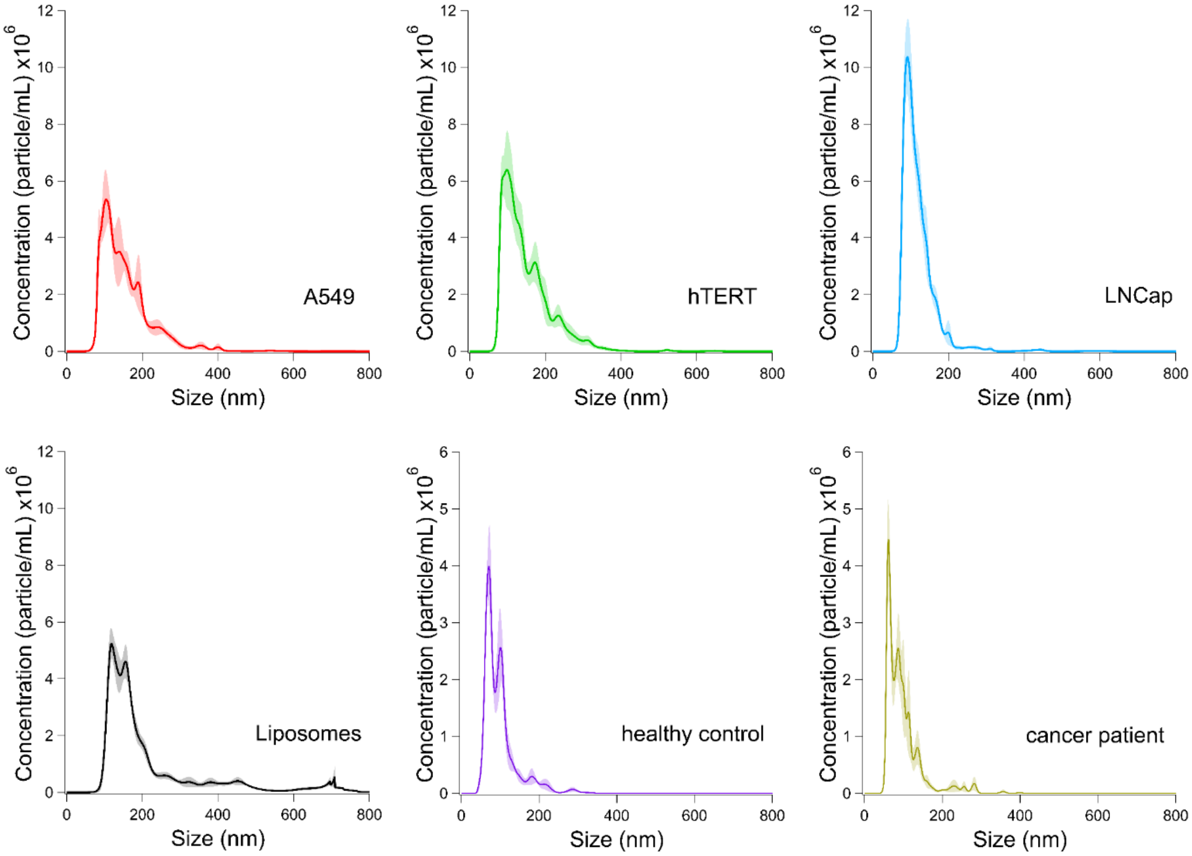


**Figure S7**. Nanoparticle tracking analysis results for commercial cell-line exosomes, liposomes, and clinically isolated exosomes. The average size for A549, hTERT, LNCap, and liposomes used for the SERS experiment were 154.6 ± 1.3, 148.9 ± 2.72, 118.6 ± 0.9, and 218.8 ± 1.5 nm, respectively. The calculated concentrations of the A549, hTERT, LNCap, and liposomes used for the SERS experiment were 2.6 x $10^9$, 2.8 x $10^9$, 2.8 x $10^9$, and 5.5 x $10^{11}$ particles/mL, respectively. The average size for clinical healthy control exosomes and cancer patient exosomes used for the SERS experiment were 106.5 ± 2.9 and 98.2 ± 2.1 nm, respectively. The calculated concentrations used for the SERS experiment were 1 x $10^9$ and 1 x $10^9$ particles/mL, respectively.

*Characterization of SERS Enhancement of Substrates*

To evaluate the SERS substrate with the high plasmonic enhancement, the spectrum of benzoic acid (BA), a Raman active molecule, was collected on the various SERS substrates. Each substrate type was tested by preparing three substrates each with 10 μL of $10^{-5}$M BA. Six spectra were collected per substrate, thus 18 spectra of BA for each substrate type were analyzed. The BA Raman signal area at 1001 ± 2.5 cm-1 was evaluated and subtracted with the Raman signal in that same region without BA. For the solid-state fabricated substrates, commercially available gold (Au) and silver (Ag) substrates were tested alongside metallic film over nanospheres (MeFON) and colloidal-state fabricated substrates, including gold nanoparticles (Au NPs), gold nanostars (S15), and silver-coated gold nanostars (S10@Ag2, S10@Ag5, S10@Ag10).

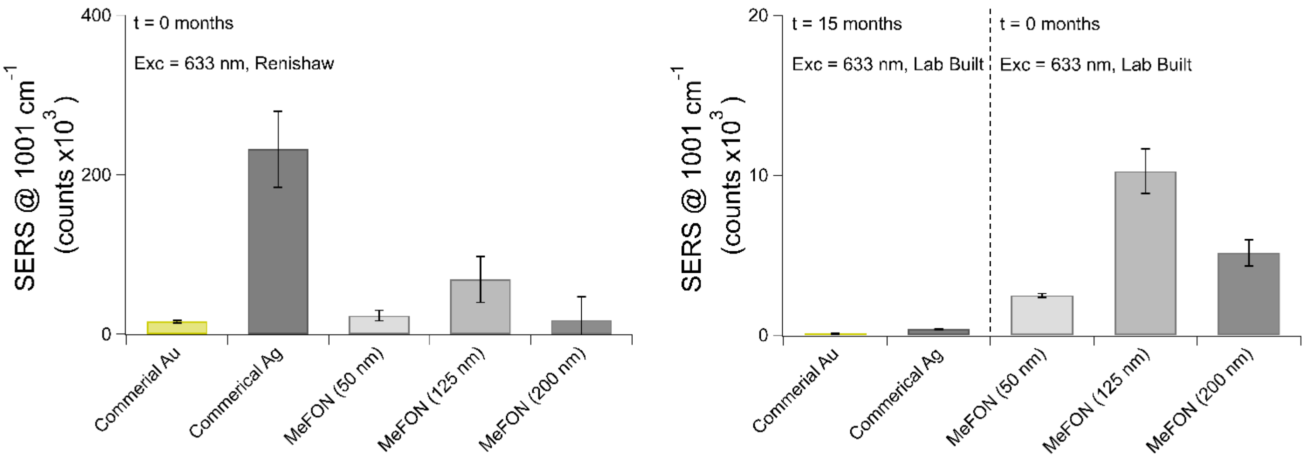


**Figure S8**. Benzoic acid's SERS intensity area at 1001 ± 2.5 $cm^{-1}$ for solid-state fabricated substrates on Renishaw and Lab Built. At 0 months (t = 0), the Ag commercial substrate had the highest SERS enhancement on a Renishaw. However, after 15 months, the commercial substrates' SERS enhancement was less compared to MeFON on a Lab Built Raman. Hence and was not further used for exosome SERS analysis.

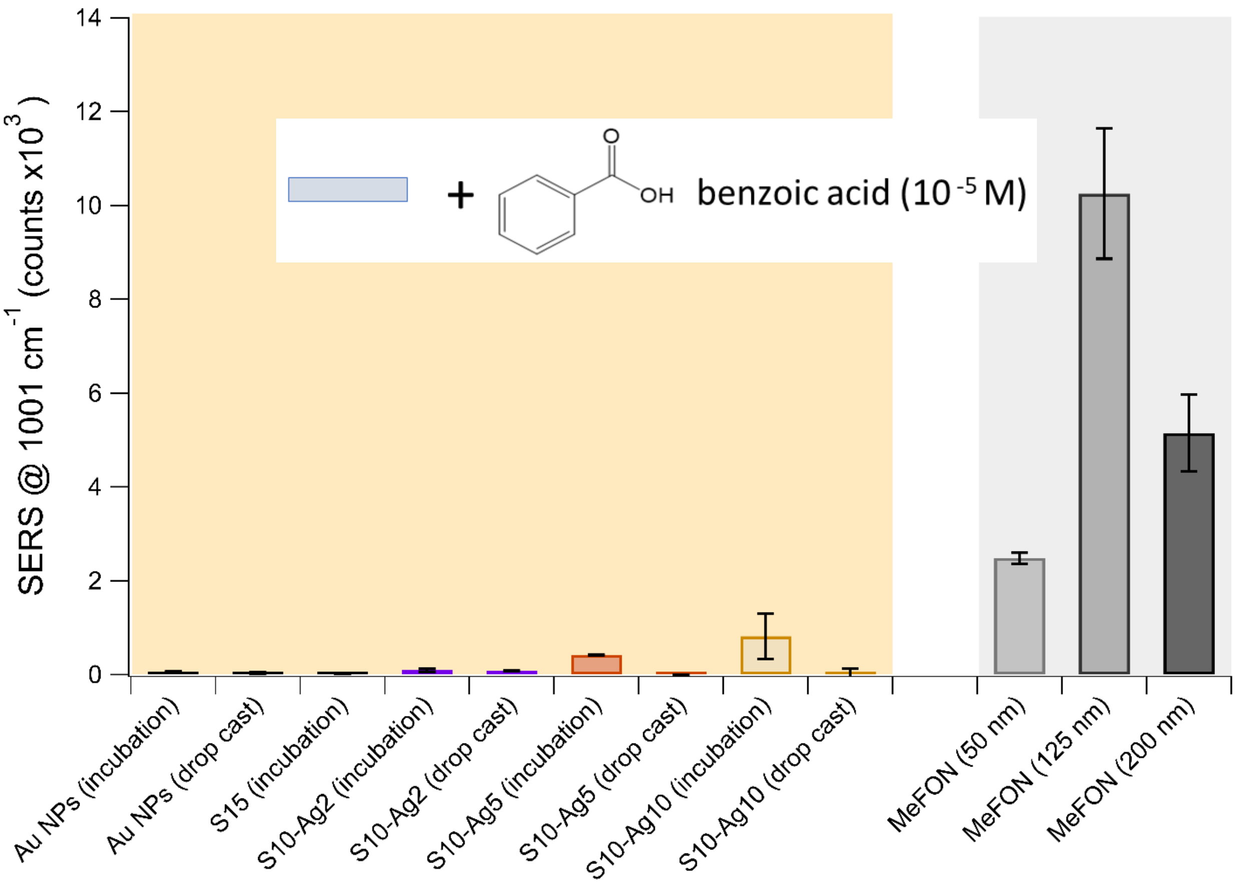


**Figure S9**. On a Lab Built Raman, Benzoic acid's SERS intensity area at 1001 ± 2.5 $cm^{-1}$ for colloidal-state (yellow region) and solid-state (gray region) fabricated substrates surveyed after blank removal. The metallic film over nanospheres (MeFON) substrates had superior SERS intensity and were used to optimize the ML methods. The MeFON of 125 nm thickness had a higher SERS enhancement. Therefore, this solid-state fabricated substrate of MeFON of 125 nm, denoted as MeFON in the main text, was further used in exosome analysis and ML model optimization. As for the colloidal-state substrates, the silver-coated gold nanostars (ASNS-Ag), specifically S10-Ag10 incubation substrates had a high SERS enhancement with a larger deviation. Therefore, the S10-Ag5 further denoted in the main text as NS, incubation substrates were used to test the optimized ML methods to examine the clinical exosomes SERS spectra.

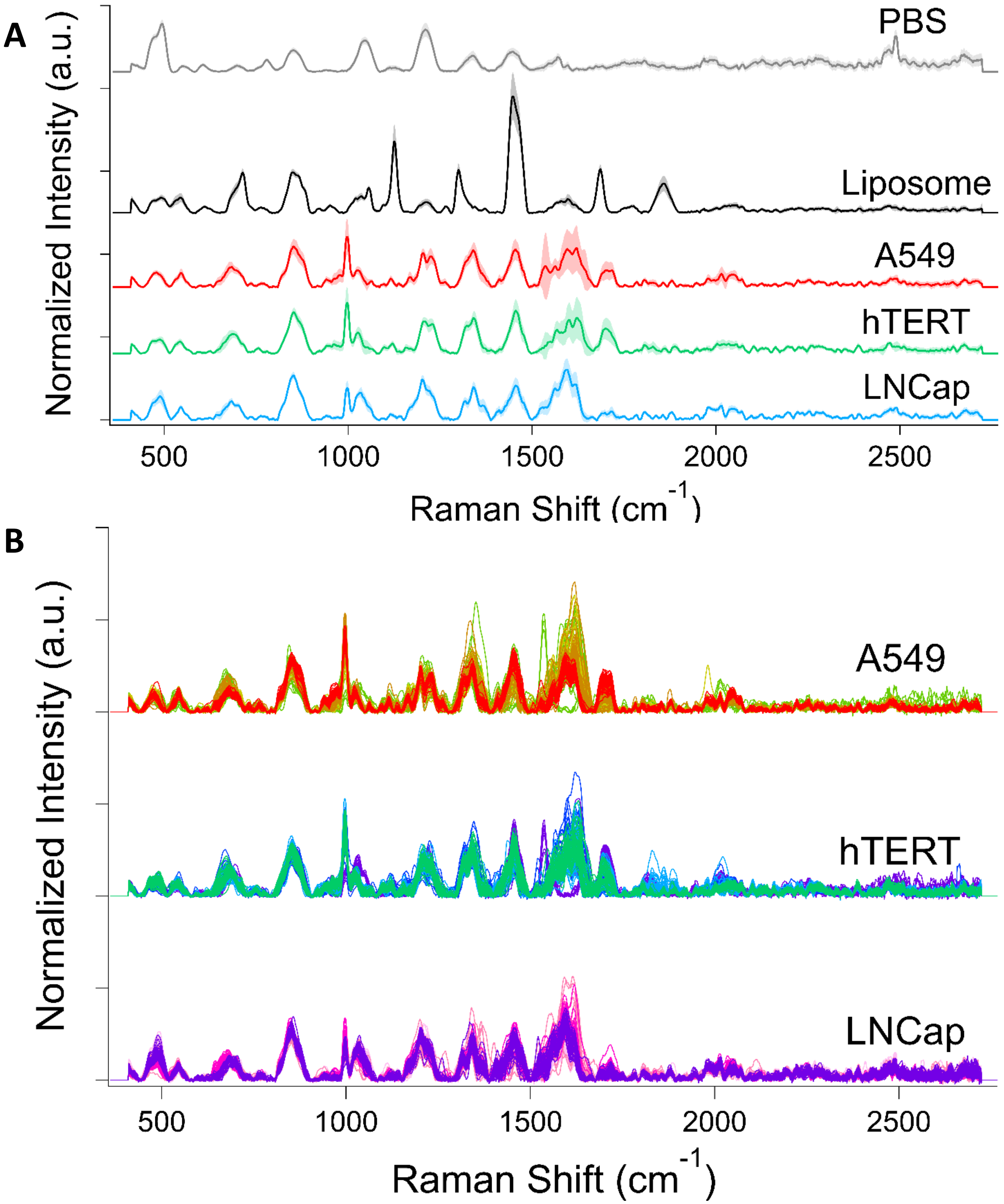


**Figure S10**. (A) Averaged normalized SERS spectra for Liposomes and PBS compared the cell-line exosomes. (B) The individual normalized spectra of A549 (N= 250), hTERT (N= 258), and LNCap (N= 205) exosomes obtained on multiple solid-state fabricated SERS substrates metallic film over nanospheres (MeFON) of 125 nm (4 substrates/ exosome). Substrates are denoted as by the different colors.

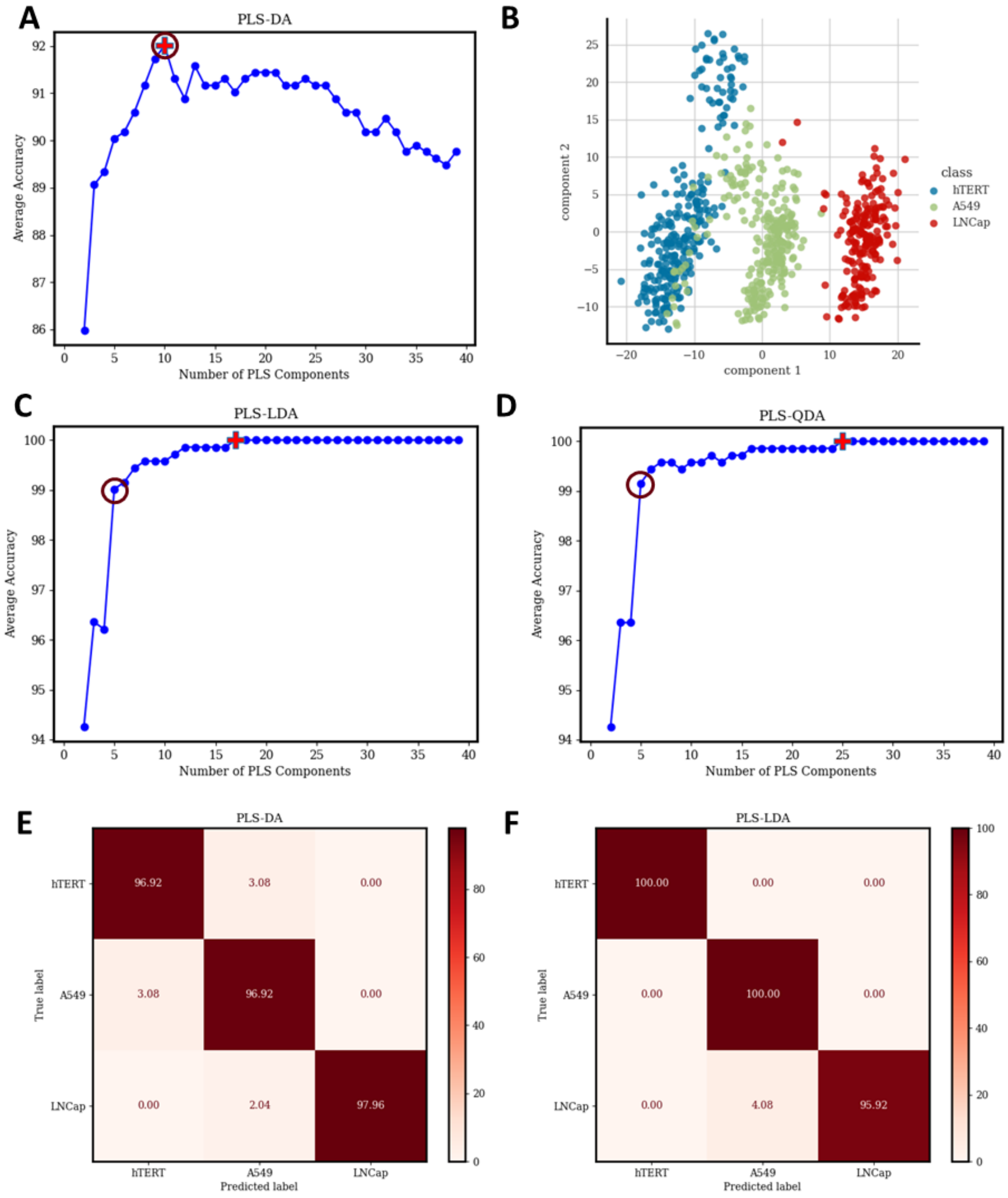


**Figure S11**. Average accuracy vs number of partial least squares (PLS) components plot for (A) discriminant analysis (DA) where the max Average accuracy is denoted as **+** and the **O** correlates to the number of components used for the classification model. For visualization, (B) the initial reduced space with the first 2 components on commercial cell-line exosomes is on the right. The hTERT exosome did have a slight cluster deviation. However, there was a clear differentiation among the three exosomes. We also explored the Average accuracy vs number of partial least squares (PLS) components plot for (C) linear (LDA) and (D) quadratic DA (QDA). E and F show the result in confusion matrix in percentage from the classification models for PLS-DA, with 10 components, and PLS-LDA, with 5 components with DA and LDA resulting in an accuracy of 92.0%, and 99.0%, respectively, which was slightly lower compared to QDA (99.2%, Figure 3) (N= 214).

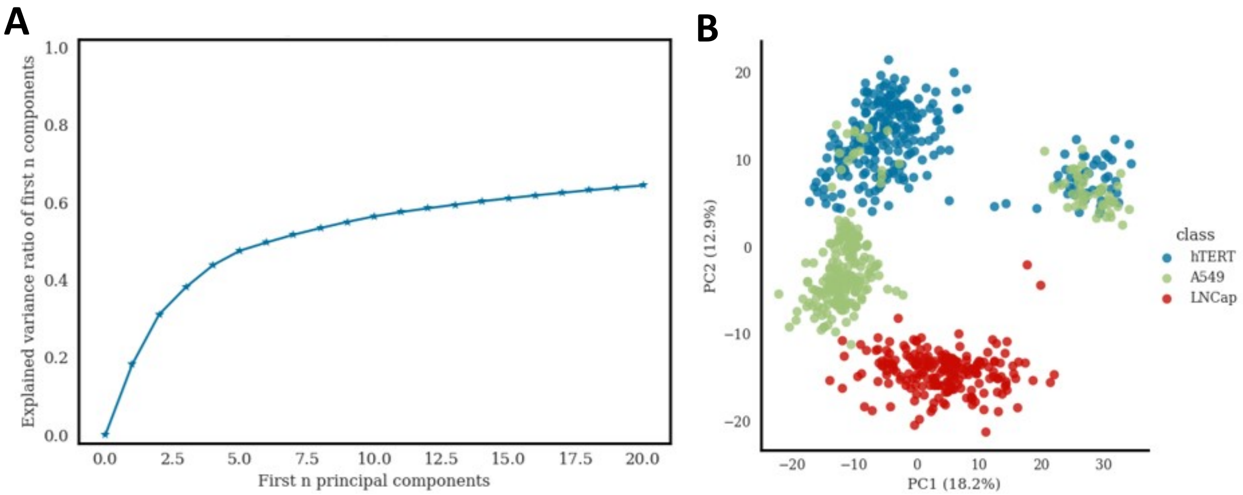


**Figure S12**. Total explained variance from the first 20 principal components from the principal component analysis (PCA) was 63% (A). For visualization, the initial reduced space with the first 2 principal components (PC1: 18.2%, PC2: 12.9% variance explained) on commercial cell-line exosomes (B). Clustering for the three exosomes was observed. However, there was a small overlap between A549 and hTERT. We suspect this is due to the CD63 protein that they both express from the ATCC report.

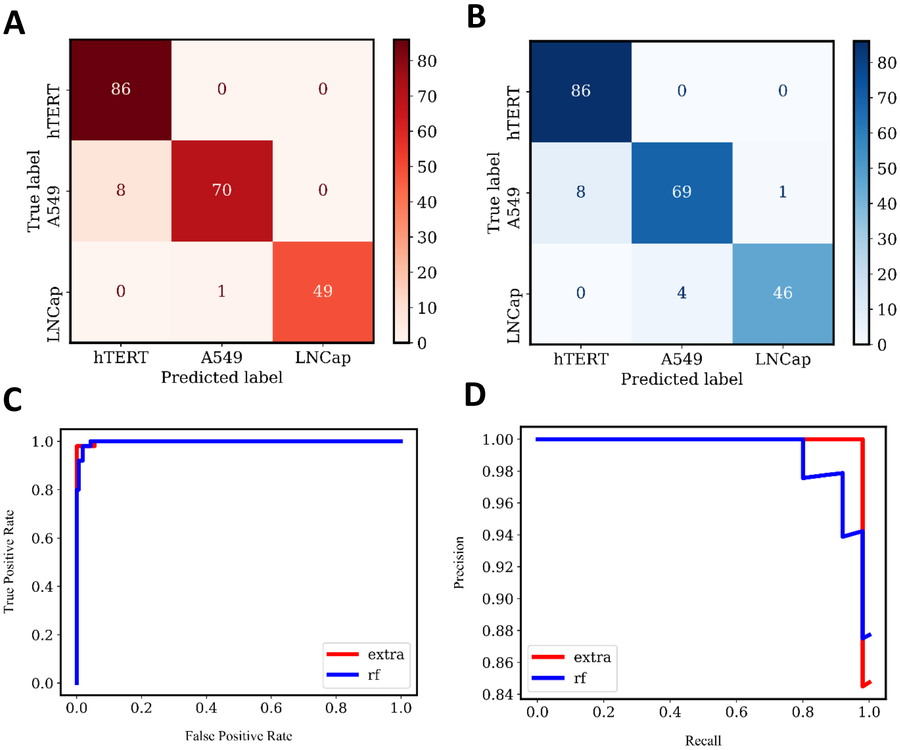


**Figure S13**. The detailed confusion matrix results of the commercial cell-line exosomes from the Extra Trees (A) and the Random Forest (B) classifiers, both with N= 214. ROC Curves (C) and Precision-Recall Curves (D) show Extra Trees (extra, red) as a better-performing classifier compared to Random Forest (rf, blue).

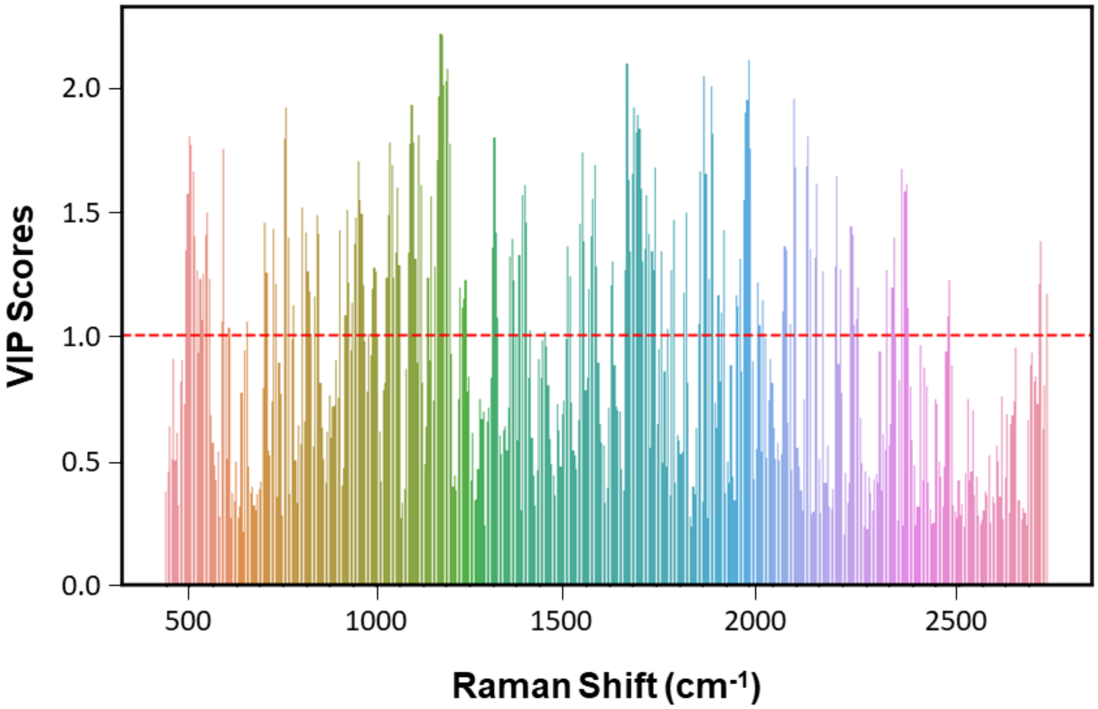


**Figure S14**. Variable Importance in Projection (VIP) scores of PLS-DA analysis on the commercial cell-line exosomes.

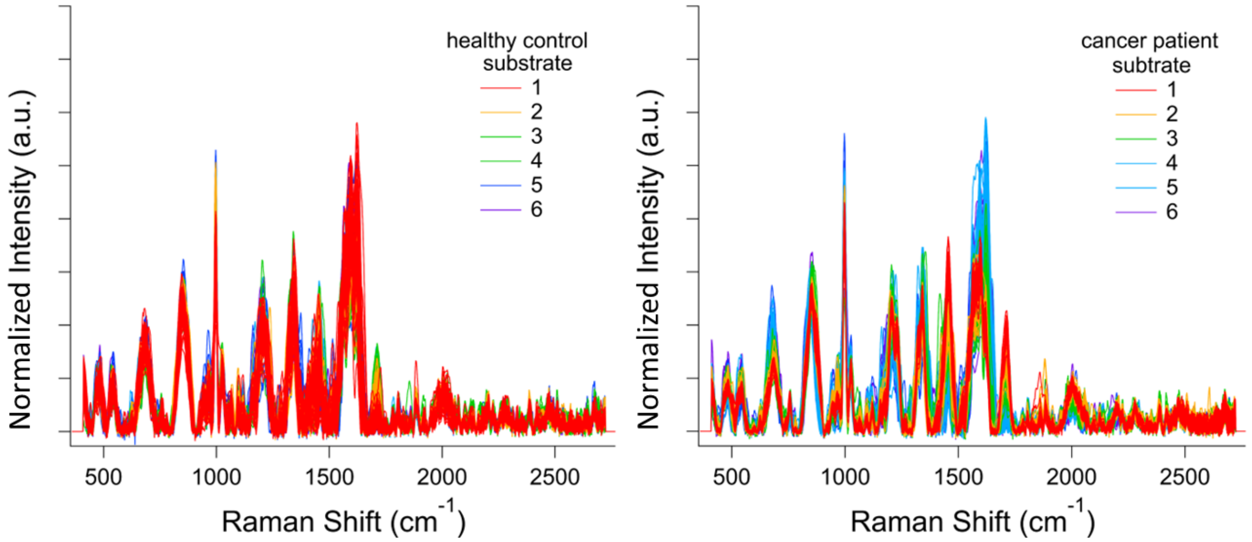


**Figure S15**. The individual normalized spectra of clinical healthy control (N= 310) and cancer patient (N= 308) exosomes obtained from multiple solid-state fabricated SERS substrates metallic film over nanospheres (MeFON) of 125 nm (6 substrates/ exosome) were used to make the averaged normalized SERS spectra for Figure 6B. Substrates are denoted as by the different colors.

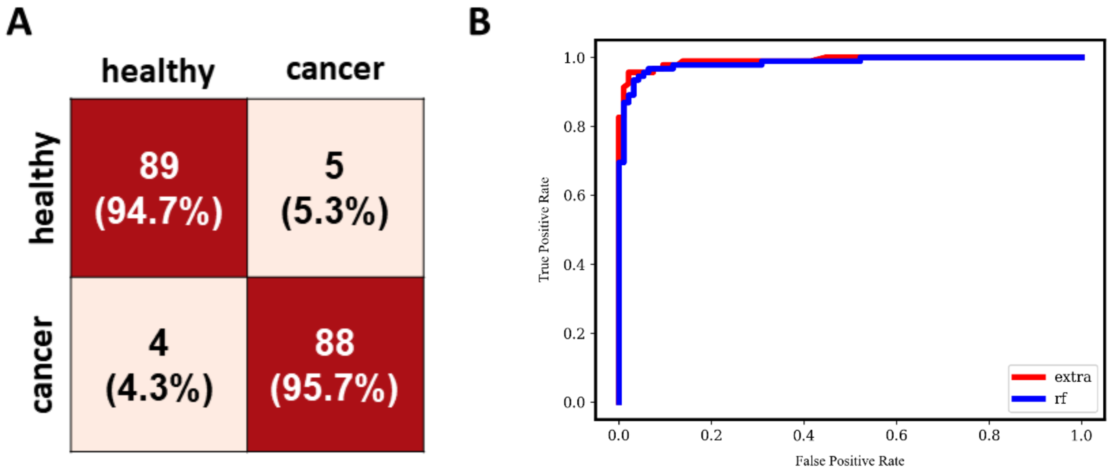


**Figure S16**. Confusion matrix (A) and ROC (B) curves of the clinical exosomes from the Random Forest, second best performing bagging model. Random Forest (rf, blue) did not outperform Extra Tree (extra, red) and had an AUC of 0.90.

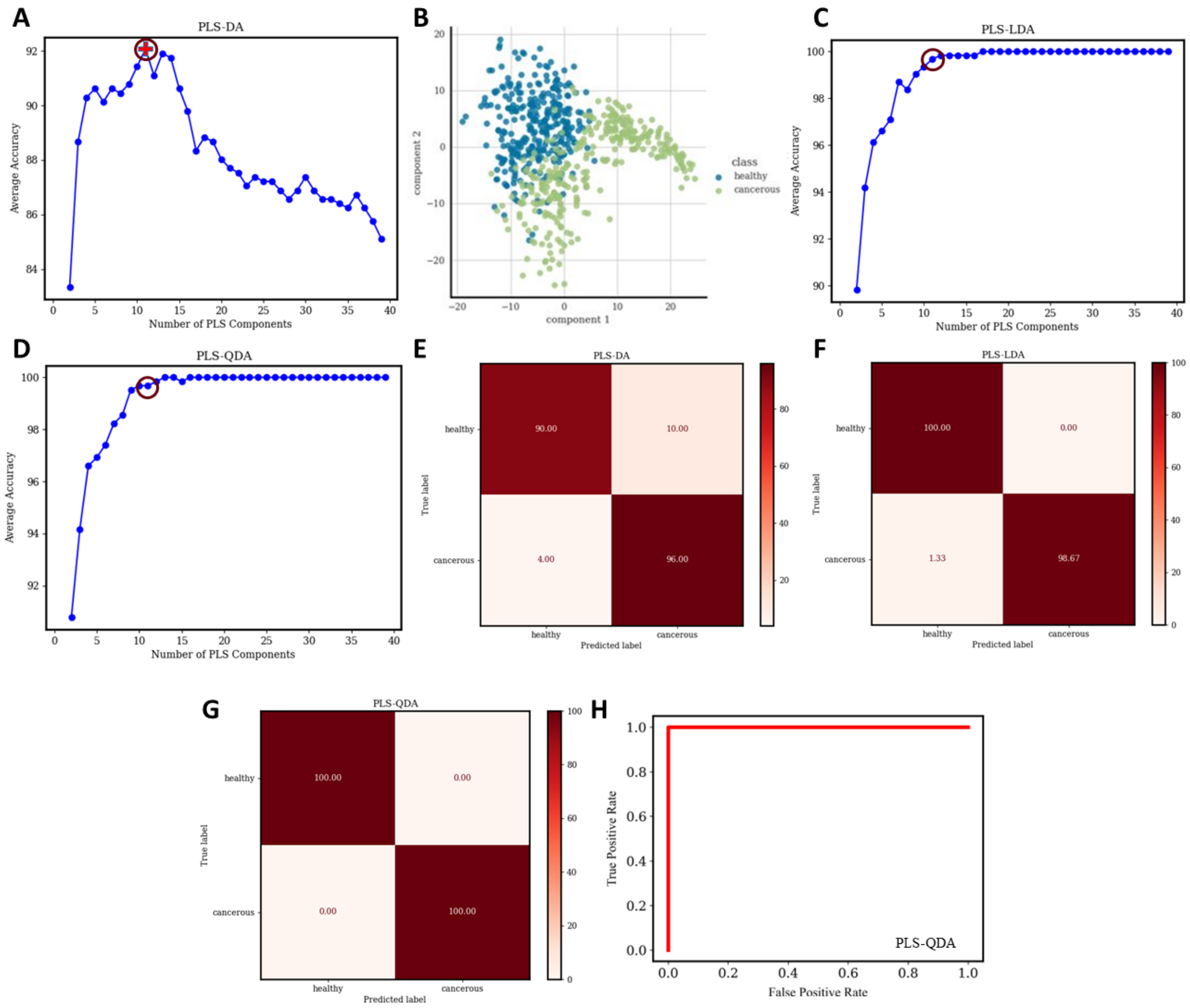


**Figure S17**. Average accuracy vs number of partial least squares (PLS) components plot for (A) discriminant analysis (DA) where the max Average accuracy is denoted as **+** and the **O** correlates to the number of components used for the classification model. For visualization, the initial reduced space with the first 2 components on commercial cell-line exosomes is in B with some overlap observed. We also explored the Average accuracy vs number of partial least squares (PLS) components plot for (C) linear (LDA) and (D) quadratic DA (QDA). E, F, and G show the result in confusion matrix in percentage from the classification models for PLS-DA, PLS-LDA, and PLS-QDA with 11 components each, with DA, LDA, and QDA resulting in an accuracy of 92.1%, 99.7% and 99.7%, respectively. The ROC curve for PLS-QDA (H) does outperform Extra Trees and had an AUC of 1.0. This suggests that the PLS/QDA could still discern changes within the two exosomes' spectra, enabling a distinction between healthy and cancerous exosomes, but could also be an indication of deceptive results due to model overfitting. Nevertheless, it's important to acknowledge that concerns persist regarding the quality of the transformed data between the clinical and commercial cell-line samples.

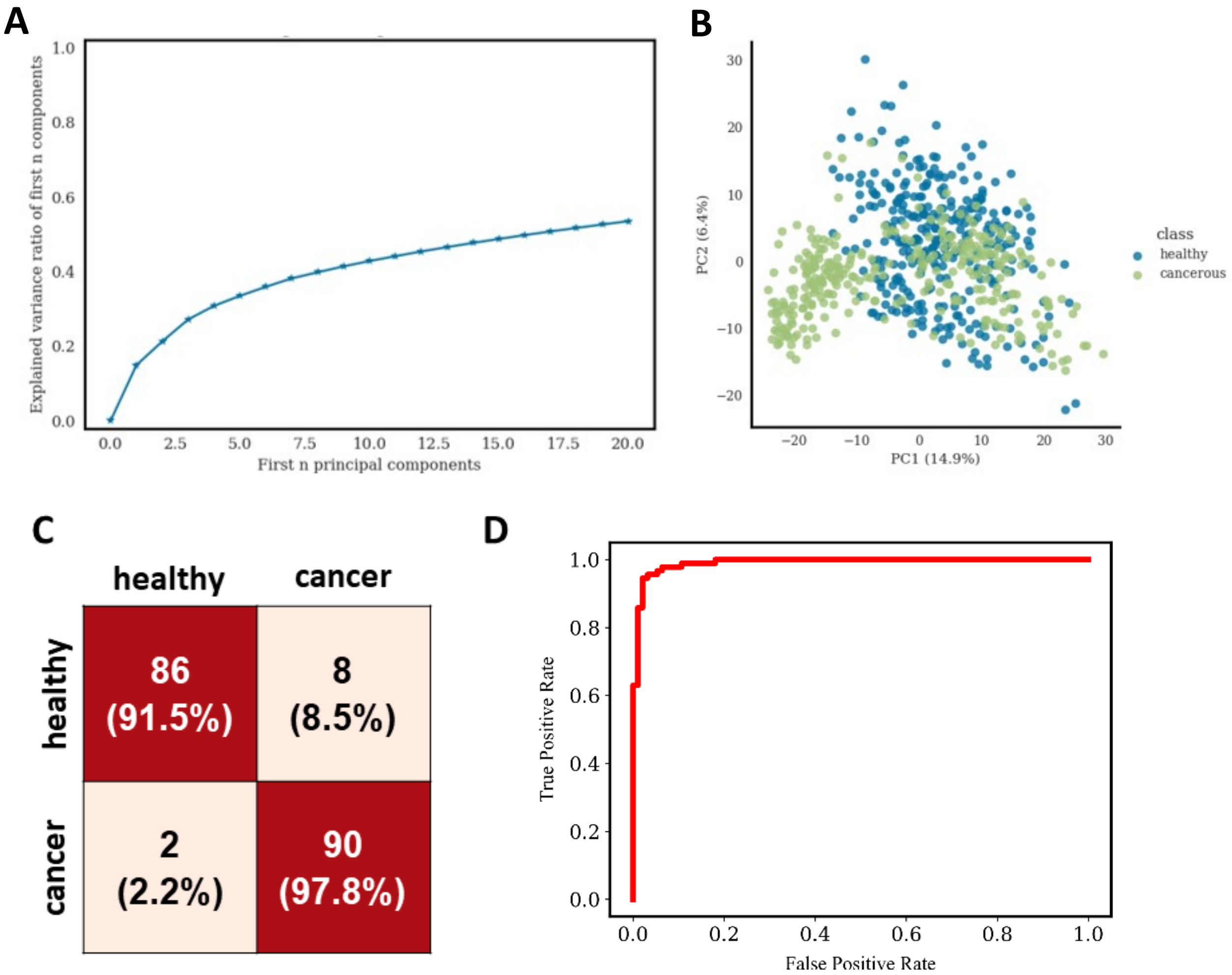


**Figure S18**. Total explained variance from the first 20 principal components from the principal component analysis (PCA) was 57% (A). For visualization, the initial reduced space with the first 2 principal components (PC1: 14.9%, PC2: 6.4% variance explained) on commercial cell-line exosomes (B). The projection displays a complete overlap between healthy and cancerous spectra, a less favorable outcome compared to the PLS space. Nevertheless, when employing SVM with various kernels, we observed model accuracies ranging from 86% to 95%, with the radial basis function kernel consistently yielding the best-performing model (Table S7) and summarized in the confusion matrix (C) and ROC curve (D). While PCA/SVM outperformed Extra Trees and had an AUC of 0.97. However, PCA/SVM explains less than 60% of the data's total variance when using 20 components, which perpetuates the concerns about this method's application on future data.

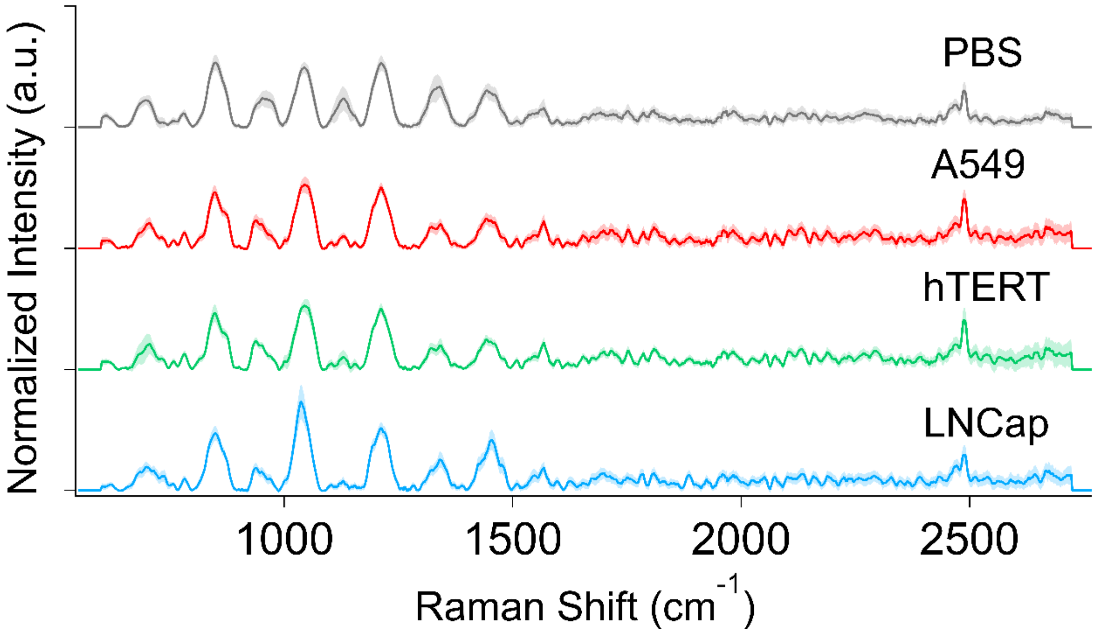


**Figure S19**. Averaged normalized SERS spectra for A549, hTERT, LNCap, and PBS on colloidal-state fabricated SERS substrates (NS of S10-Ag5 incubated on silicon).

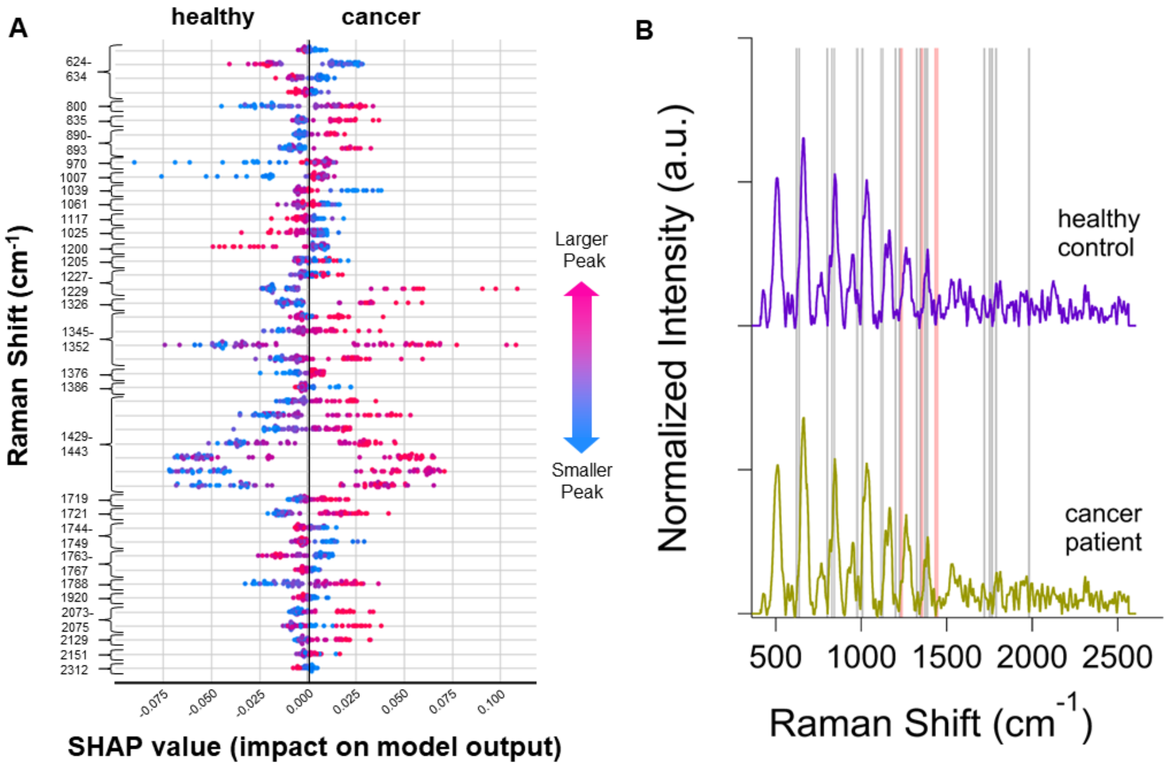


**Figure S20**. SHAP beeswarm plot shows the top 5% of the most important Raman Shifts in the classification of the clinical exosomes on the colloidal nanostars silicon substrate.

**Table S1**. Initial classification analysis was performed on full cell-culture spectra (984 peaks) using the solid-state substrate. 15 classifiers were compared using the average accuracy, area under the curve, recall, precision, F1-score, kappa, and MCC of 10 cross-validation sets. The top 5 classifiers (bolded), based on AUC, were selected for continual testing and optimization. The random state was set at a value of 101.

| Model | Accuracy | AUC | Recall | Precision | F1 | Kappa | MCC |
|---|---|---|---|---|---|---|---|
| **Extra Trees Classifier** | 0.972 | 0.999 | 0.972 | 0.974 | 0.972 | 0.958 | 0.959 |
| **Light Gradient Boosting Machine** | 0.974 | 0.999 | 0.974 | 0.976 | 0.974 | 0.961 | 0.962 |
| **CatBoost Classifier** | 0.978 | 0.998 | 0.978 | 0.980 | 0.978 | 0.967 | 0.968 |
| **Random Forest Classifier** | 0.974 | 0.998 | 0.974 | 0.976 | 0.974 | 0.961 | 0.962 |
| **Gradient Boosting Classifier** | 0.964 | 0.997 | 0.964 | 0.968 | 0.964 | 0.946 | 0.947 |
| K Neighbors Classifier | 0.974 | 0.996 | 0.974 | 0.976 | 0.974 | 0.961 | 0.932 |
| Linear Discriminant Analysis | 0.956 | 0.989 | 0.956 | 0.958 | 0.956 | 0.933 | 0.935 |
| Ada Boost Classifier | 0.918 | 0.987 | 0.918 | 0.925 | 0.918 | 0.876 | 0.879 |
| Naive Bayes | 0.932 | 0.977 | 0.932 | 0.937 | 0.931 | 0.897 | 0.900 |
| Logistic Regression | 0.363 | 0.952 | 0.363 | 0.132 | 0.193 | 0.000 | 0.000 |
| Decision Tree Classifier | 0.906 | 0.928 | 0.906 | 0.910 | 0.906 | 0.858 | 0.860 |
| Quadratic Discriminant Analysis | 0.433 | 0.570 | 0.433 | 0.431 | 0.427 | 0.141 | 0.143 |
| Dummy Classifier | 0.363 | 0.500 | 0.363 | 0.132 | 0.193 | 0.000 | 0.000 |
| SVM - Linear Kernel | 0.443 | 0.000 | 0.443 | 0.309 | 0.304 | 0.157 | 0.207 |
| Ridge Classifier | 0.415 | 0.000 | 0.415 | 0.415 | 0.291 | 0.083 | 0.176 |

**Table S2**. PLS/DA results of the commercial cell-line exosomes on the MeFON substrate.

| DA | Accuracy | AUC | Recall | Precision | F1 |
|---|---|---|---|---|---|
| DA | 0.920 | 0.920 | 0.925 | 0.931 | 0.925 |
| linear | 0.990 | 1.000 | 0.989 | 0.991 | 0.990 |
| quadratic | 0.992 | 0.991 | 0.991 | 0.992 | 0.992 |

**Table S3**. PCA/SVM results of the commercial cell-line exosomes on the MeFON substrate.

| SVM | Accuracy | AUC | Recall | Precision | F1 |
|---|---|---|---|---|---|
| linear | 0.962 | 0.000 | 0.962 | 0.963 | 0.962 |
| polynomial | 0.972 | 0.000 | 0.972 | 0.974 | 0.972 |
| radial basis function | 0.972 | 0.000 | 0.972 | 0.974 | 0.972 |

**Table S4**. Optimization parameters that differ from the default settings for top classifiers.

| Model | Parameter | Value |
|---|---|---|
| Random Forest Classifier (scikit-learn 1.2.1) | estimators | 170 |
| | max_depth | 8 |
| | min_sample_leaf | 2 |
| | max_features | log2 |
| | bootstrap | False |
| Gradient Boosting Classifier (scikit-learn 1.2.1) | learning_rate | 0.5 |
| | n_estimators | 270 |
| | subsample | 0.85 |
| | min_samples_split | 10 |
| | max_features | sqrt |
| LGBM Classifier (v. 3.3.5) | n_leaves | 31 |
| | learning_rate | 0.05 |
| | n_estimators | 160 |
| | min_split_gain | 0.8 |
| | min_child_samples | 61 |
| | reg_alpha | 0.0005 |
| | reg_lambda | 0.005 |
| Extra Trees Classifier (scikit-learn 1.2.1) | n_jobs | -1 |

| Model | Parameter | Value |
|---|---|---|
| CatBoost Classifier (v. 1.2) | nan_mode | min |
| | gpu_ram_part | 0.95 |
| | eval_metric | Multiclass |
| | iterations | 80 |
| | leaf_estimation_method | Newton |
| | grow_policy | Symmetric Tree |
| | penalities_coefficient | 1 |
| | boosting_type | Plain |
| | feature_boarder_type | GreedyLog Sum |
| | devices | -1 |
| | l2_leaf_ref | 1 |
| | random_strength | 0.2 |
| | rsm | 1 |
| | model_size_reg | 0.5 |
| | depth | 6 |
| | border_count | 128 |
| | leaf_estimation_backtracking | Any Improvement |
| | min_data_in_leaf | 1 |
| | loss_function | MultiClass |
| | learning_rate | 0.5 |
| | score_function | Cosine |
| | leaf_estimation_iterations | 1 |
| | bootstrap_type | Bayesian |
| | max_leaves | 64 |

**Table S5**. The top classifiers' performance after using SHAP to construct a model using only the top 5% most important spectral pixel, consisting of 49 pixels.

| Model | Accuracy | AUC | Recall | Precision | F1 | Kappa | MCC |
|---|---|---|---|---|---|---|---|
| Random Forest | 0.9619 | 0.9968 | 0.9619 | 0.9641 | 0.9617 | 0.9425 | 0.9438 |
| Gradient Boosting | 0.9839 | 0.9973 | 0.9839 | 0.9848 | 0.9839 | 0.9757 | 0.9762 |
| Light Gradient Boosting Machine | 0.9679 | 0.9963 | 0.9679 | 0.9695 | 0.9679 | 0.9516 | 0.9524 |
| Extra Trees | 0.9699 | 0.9967 | 0.9699 | 0.9723 | 0.9698 | 0.9546 | 0.956 |
| CatBoost | 0.9739 | 0.9978 | 0.9739 | 0.9764 | 0.9739 | 0.9606 | 0.962 |
| K-neighbor | 0.9699 | 0.987 | 0.9699 | 0.9714 | 0.9698 | 0.9546 | 0.9554 |

Table S6. PLS/DA results of the clinical exosomes on the MeFON substrate.

| DA | Accuracy | AUC | Recall | Precision | F1 |
|---|---|---|---|---|---|
| DA | 0.921 | 0.922 | 0.614 | 0.615 | 0.614 |
| linear | 0.997 | 0.996 | 0.997 | 0.997 | 0.997 |
| quadratic | 0.997 | 0.997 | 0.997 | 0.997 | 0.997 |

Table S7. PCA/SVM results of the clinical exosomes on the MeFON substrate.

| SVM | Accuracy | AUC | Recall | Precision | F1 |
|---|---|---|---|---|---|
| linear | 0.863 | 0.000 | 0.857 | 0.869 | 0.862 |
| polynomial | 0.912 | 0.000 | 0.842 | 0.980 | 0.904 |
| radial basis function | 0.947 | 0.000 | 0.921 | 0.973 | 0.946 |
